# A knockoff calibration method to avoid over-clustering in single-cell RNA-sequencing

**DOI:** 10.1101/2024.03.08.584180

**Authors:** Alan DenAdel, Michelle L. Ramseier, Andrew W. Navia, Alex K. Shalek, Srivatsan Raghavan, Peter S. Winter, Ava P. Amini, Lorin Crawford

## Abstract

Standard single-cell RNA-sequencing (scRNA-seq) pipelines nearly always include unsupervised clustering as a key step in identifying biologically distinct cell types. A follow-up step in these pipelines is to test for differential expression between the identified clusters. When algorithms over-cluster, downstream analyses will produce inflated *P* -values resulting in increased false discoveries. In this work, we present `callback` (**Cal**i**b**r**a**ted **C**lustering via **K**nockoffs): a new method for protecting against over-clustering by controlling for the impact of reusing the same data twice when performing differential expression analysis, commonly known as “double-dipping”. Importantly, our approach can be applied to a wide range of clustering algorithms. Using real and simulated data, we show that `callback` provides state-of-the-art clustering performance and can rapidly analyze large-scale scRNA-seq studies, even on a personal laptop.

## Main

Recent advances in single-cell RNA sequencing (scRNA-seq) technologies have enabled the generation of datasets that contain the transcriptomic profiles of thousands to millions of individual cells [1, 2]. Unless an additional assay is paired with sequencing (e.g., CITE-seq [3]), cell type labels are not provided with the corresponding genomic profiles. This has led to many scRNA-seq bioinformatic pipelines requiring both (i) clustering to identify putative cell types based on shared gene expression covariation and (ii) differential gene expression analysis between cells in each cluster to identify “marker genes” uniquely expressed by each putative cell type. The most commonly used software packages, such as `Seurat` [4] and `Scanpy` [5], perform these two steps on the same dataset. This double use of data is often referred to as “circular analysis” or “double-dipping,” and is known to result in highly inflated *P* -values, even in the null case when gene expression is identically distributed and there are no true groupings that distinguish cell populations [6, 7]. Due to the miscalibrated test statistics produced by circular analyses, it is challenging to assess whether the genes found to be differentially expressed between two putative cell groups are “real” or solely identified due to chance based on the way that the cells are being partitioned by the clustering algorithm that is being used. Importantly, simple solutions such as sample splitting between cells do not appropriately correct for this type of post-selective inference [7].

Several methods have been recently developed to correct for post-selective inference after clustering. These methods include: (i) an approximate test based on the truncated normal distribution [8], (ii) a data splitting strategy that splits data at the level of individual gene counts [7], and (iii) using synthetic null variables called knockoffs for calibrating hypothesis testing [6]. The point of each of these methods is to identify an appropriate hypothesis testing significance threshold to account for the statistical inflation that occurs due to the double use of data. However, none of these tests inform if (or how) the re-clustering of cells should be done. They simply return a list of calibrated *P* -values. As a result, approaches for protecting against over-clustering have recently been proposed including “single cell significance of hierarchical clustering” (`sc-SHC`) and “clustering hierarchy optimization by iterative random forests” (`CHOIR`)[9, 10]. Here, we introduce `callback` (**Cal**i**b**r**a**ted **C**lustering via **K**nockoffs): a method that integrates the negative control variable framework of knockoffs [11, 12] to the problem of identifying the number of clusters that have statistical support in a single-cell dataset. Our approach can be paired with any existing clustering algorithm that has a hyperparameter for tuning the number of clusters and it makes no strong assumptions about the input data. We statistically motivate the need for an algorithm like `callback`, evaluate its utility against other recently proposed clustering correction methods, and demonstrate its ability to efficiently scale to large-scale scRNA-seq studies.

The `callback` algorithm consists of three simple steps (Methods). First, we generate synthetic null variables, formally called knockoff features [11], where we augment the single-cell data being analyzed with “fake” genes that are known not to contribute to any unique cell type but that match the real data in distribution. Second, we perform both preprocessing and clustering on this augmented dataset. Third, we calibrate the number of inferred clusters by using a hypothesis testing strategy with a data-dependent threshold to determine if there is a statistically significant difference between groups and if re-clustering should occur (Fig. 1a). The synthetic knockoff genes act as negative control variables; they go through the same analytic steps as the real data and are presented with the same opportunity to be identified as marker genes. The `callback` algorithm uses the guiding principle that well-calibrated clusters (i.e., those representing real groups) should have statistically significant differentially expressed genes after correcting for post-selective testing, while over-clustered groups will have greatly fewer. We use this rule to iteratively re-cluster cells until the inferred clusters are well-calibrated and the observed differences in expression between groups are not due to the effects of double-dipping.

**Figure 1.**
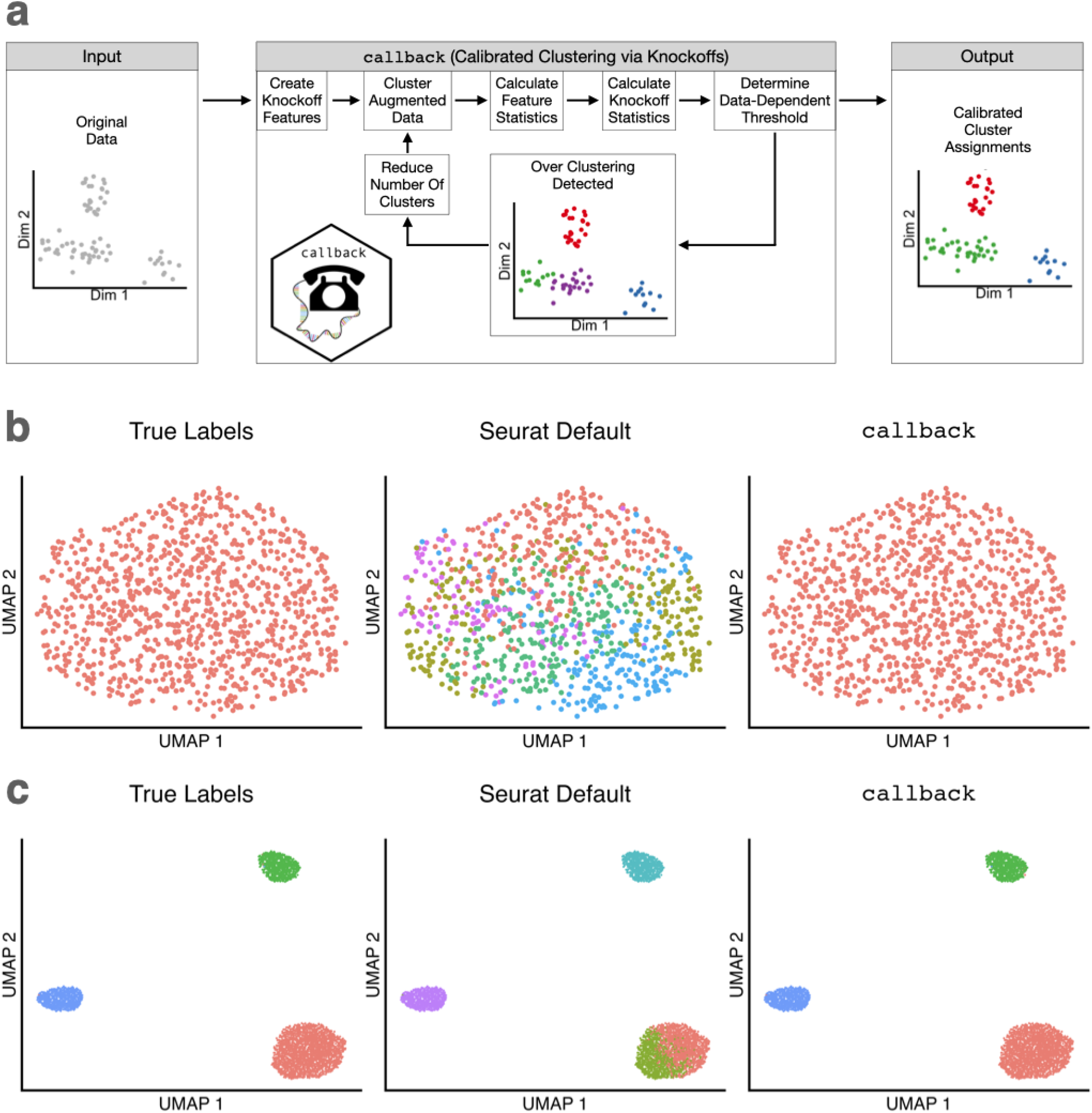
Overview of the callback algorithm and examples of results from different clustering approaches on simple simulated datasets. **(a)** Schematic of the clustering workflow with the `callback` approach. **(b)** Demonstration of the traditional clustering framework versus the alternative using `callback` for simulated data with one known group. Panels left to right show the true labels, clusters found using the Louvain algorithm with default parameter settings in `Seurat`, and the clusters found using the same Louvain algorithm paired with `callback`. **(c)** Demonstration of the traditional clustering framework versus the alternative using `callback` for simulated data with three known groups. Panels left to right show the true labels, clusters found using the Louvain algorithm with default parameter settings in `Seurat`, and the clusters found using the same Louvain algorithm paired with `callback`.

As a simple proof-of-concept, we simulated single-cell gene expression data to compare the clusters found by the widely used Louvain algorithm with default parameter settings in `Seurat` (with the `FindClusters` function where the resolution parameter is set to 0.8) versus using the same Louvain algorithm paired with `callback`. We generated data under two scenarios. In the first scenario, there was only one true “cell type”. Here, the default approach with `Seurat` incorrectly identified four clusters while `callback` correctly identified only a single cluster (Fig. 1b). In the second scenario, we simulated the data such that there were three true cell types. In this case, the `Seurat` default incorrectly identified four clusters by splitting the larger group into two clusters whereas `callback` correctly identified three clusters (Fig. 1c).

To evaluate the performance of `callback` on real single-cell RNA sequencing studies, we analyzed 20 different tissues from the Tabula Muris dataset [13]. We compared `callback` with two recently proposed methods for preventing over-clustering: (i) single-cell significance of hierarchical clustering (`sc-SHC`) [9] and (ii) clustering hierarchy optimization by iterative random forests (`CHOIR`) [10]. Both of these methods utilize hierarchical clustering paired with permutation tests to decide whether or not to merge clusters.

All `callback` results are determined using the Louvain algorithm. We analyzed the 20 different tissues separately and evaluated the performance of each method by comparing their inferred cluster assignments to the manually curated cell type annotations from the original Tabula Muris study. To empirically assess the relative quality of clustering assignments, we utilized common metrics including the adjusted Rand index (ARI), the Jaccard index, the Fowlkes-Mallows index (FMI), *V* -measure, completeness, and homogeneity [14]. We include a vignette on these cluster evaluation metrics showing their behavior in a simple case study of over-clustering and under-clustering (Supplementary Note and Fig. S1). In the main text, we focus on ARI due to its popularity in the literature [14] and *V* -measure because it is the harmonic mean of completeness and homogeneity and balances the impact of over-clustering and under-clustering (Supplementary Note).

When evaluated by ARI (Fig. 2a), *V* -measure (Fig. 2b), completeness (Fig. S2), homogeneity (Fig. S3), Jaccard index (Fig. S4), and FMI (Fig. S5), `callback` shows state-of-the-art performance. In particular, when evaluated by ARI, `callback` performs best in 17 out of the 20 tissues, `sc-SHC` performs best in 2 tissues, and `CHOIR` performs best in 1 tissue. Similarly, when evaluated by *V* -measure, `callback` performs best in 18 tissues, while `sc-SHC` and `CHOIR` perform best in 1 tissue each. The clustering results for all algorithms across all 20 tissues are displayed via uniform manifold approximation and projection (UMAP) plots in Figs. S6-S25 (for visualization purposes only). For many tissues, `CHOIR` tended to group cells into many small sub-populations; while, for other tissues, `sc-SHC` severely under-clustered and failed to find any distinct cell types at all, returning only a single group (e.g., aorta, brain myeloid, and pancreas). In the diaphragm tissue, which contains five manually curated cell types, `callback` and `sc-SHC` matched the five manually curated cell type labels almost exactly, while `CHOIR` seemingly over-clustered the data (Fig. 2c). On the other hand, in the limb muscle dataset, which contains six manually curated cell types, `callback` finds six clusters that closely match the manually curated labels (ARI = 0.97 and *V* –measure = 0.95), while `sc-SHC` finds 8 clusters (ARI = 0.74 and *V* -measure = 0.79), and `CHOIR` finds 16 clusters (ARI = 0.40 and *V* -measure = 0.69) (Fig. 2d). Importantly, `callback` exhibited better computational efficiency (i.e., shorter runtime) than the other methods. While implementing each method on a personal laptop with 6 cores, `callback` was overall the fastest, `sc-SHC` exhibited a similarly short runtime, and `CHOIR` was the slowest (Fig. 2e). For example, in the fat tissue, `callback` finished 1 minute faster than `sc-SHC` and 15.6 minutes faster than `CHOIR`.

**Figure 2.**
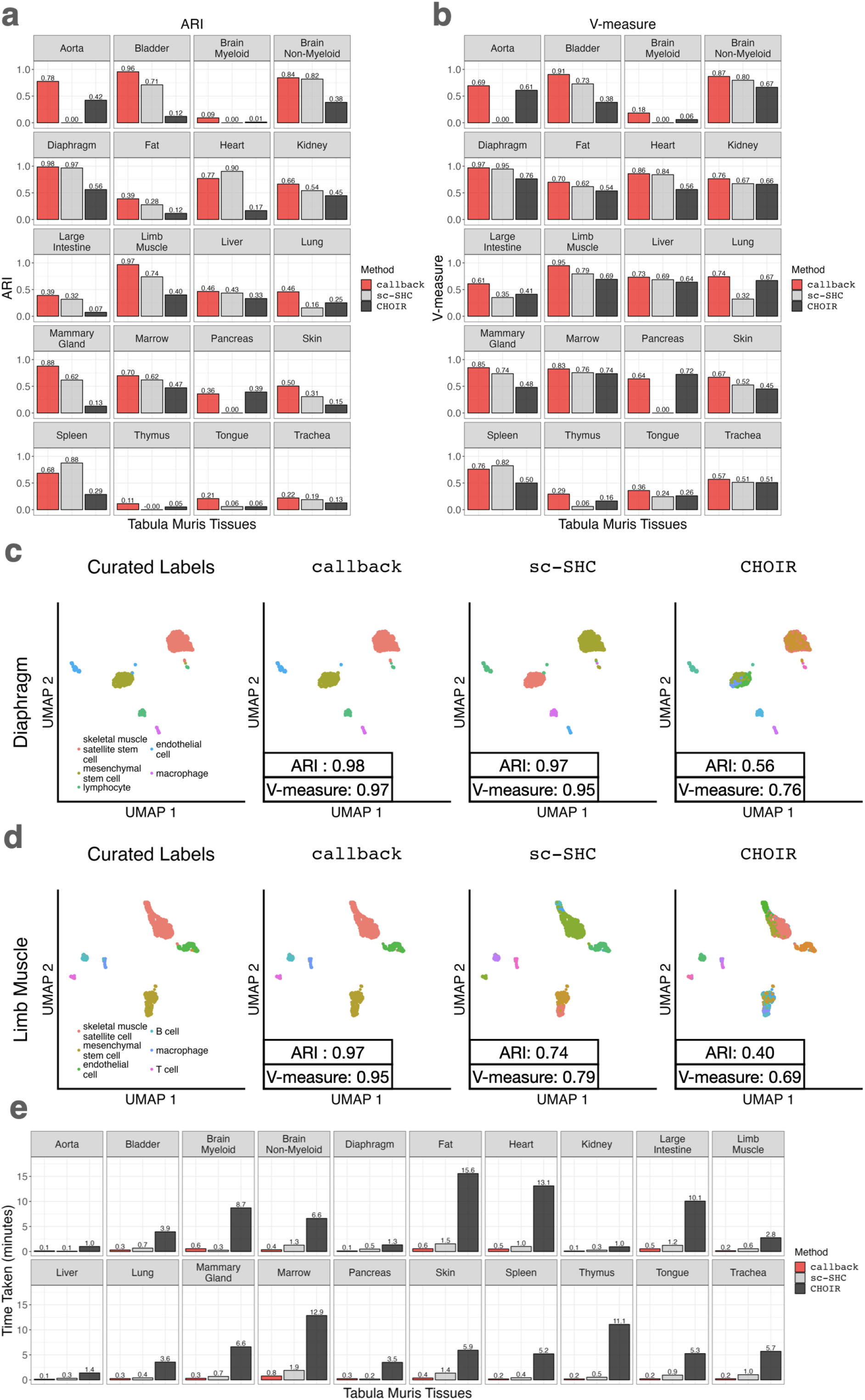
The callback algorithm shows state-of-the-art performance according to commonly used cluster quality metrics when compared to competing methods in the Tabula Muris dataset. **(a-b)** Comparison of `callback`, `sc-SHC`, and `CHOIR` using **(a)** ARI and **(b)** *V* –measure for each tissue. **(c-d)** Uniform manifold approximation and projection (UMAP) plots displaying the cell type annotations for **(c)** the diaphragm tissue and **(d)** the limb muscle tissue datasets, respectively. From left to right, we show the manually curated labels from the original study and clusters inferred by `callback`, `sc-SHC`, and `CHOIR`, respectively. **(e)** Runtime comparison of `callback`, `sc-SHC`, and `CHOIR` for each tissue in the Tabula Muris dataset. Each method was run using 6 cores on a personal laptop.

In order to show that `callback` generates useful hypotheses for downstream analyses, we further compared the clusters determined by the default `Seurat` implementation of the Louvain algorithm to the clusters determined by using the Louvain algorithm with `callback` for the limb muscle tissue in the Tabula Muris study (Fig. 3a-c). Using the `FindMarkers` function in Seurat, we identified the top 10 marker genes for each inferred cluster from both approaches. Qualitatively, the default Louvain implementation appears over-clustered, where inferred clusters 1, 2, 6, and 7 show similar marker gene expression to one another, as do inferred clusters 3 and 5 (Fig. 3d). In contrast, the groups found by `callback` show much less shared expression between clusters (Fig. 3e). To further investigate whether cells had been over-clustered by the default Louvain algorithm, we performed differential expression analysis between its inferred clusters and observed a high correlation in *P* -values when comparing (i) inferred clusters 1 and 2 versus 3 (Pearson correlation *r* = 0.923) and (ii) inferred clusters 1 and 2 versus 5 (*r* = 0.925) (Fig. 3f). For the default Louvain algorithm, the inferred clusters 1 and 2 both correspond to skeletal muscle satellite cells as annotated by the Tabula Muris Consortium, and inferred clusters 3 and 5 correspond to mesenchymal stem cells. As a comparison, only the inferred clusters 1 and 2 from `callback` correspond to skeletal muscle satellite and mesenchymal stem cells, respectively. Differential expression analysis for the `callback` clusters (Fig. 3g) results in 506 differentially expressed genes (adjusted *P* -value *<* 0.05 and an absolute log-fold change greater than one) which include many known skeletal muscle satellite cell markers up-regulated in the inferred cluster 1 relative to the inferred cluster 2 (e.g., *Des, Chodl, Myl12a, Asb5, Sdc4, Apoe, Musk, Myf5, Chrdl2, Notch3*) [15] and mesenchymal stem cell type markers up-regulated in the inferred cluster 2 relative to the inferred cluster 1 (e.g., *Col6a3, Col1a1, Igfbp6, Pdgfra, C1s, Mfap5, Ecm1, Dcn, Dpep1*) [16].

**Figure 3.**
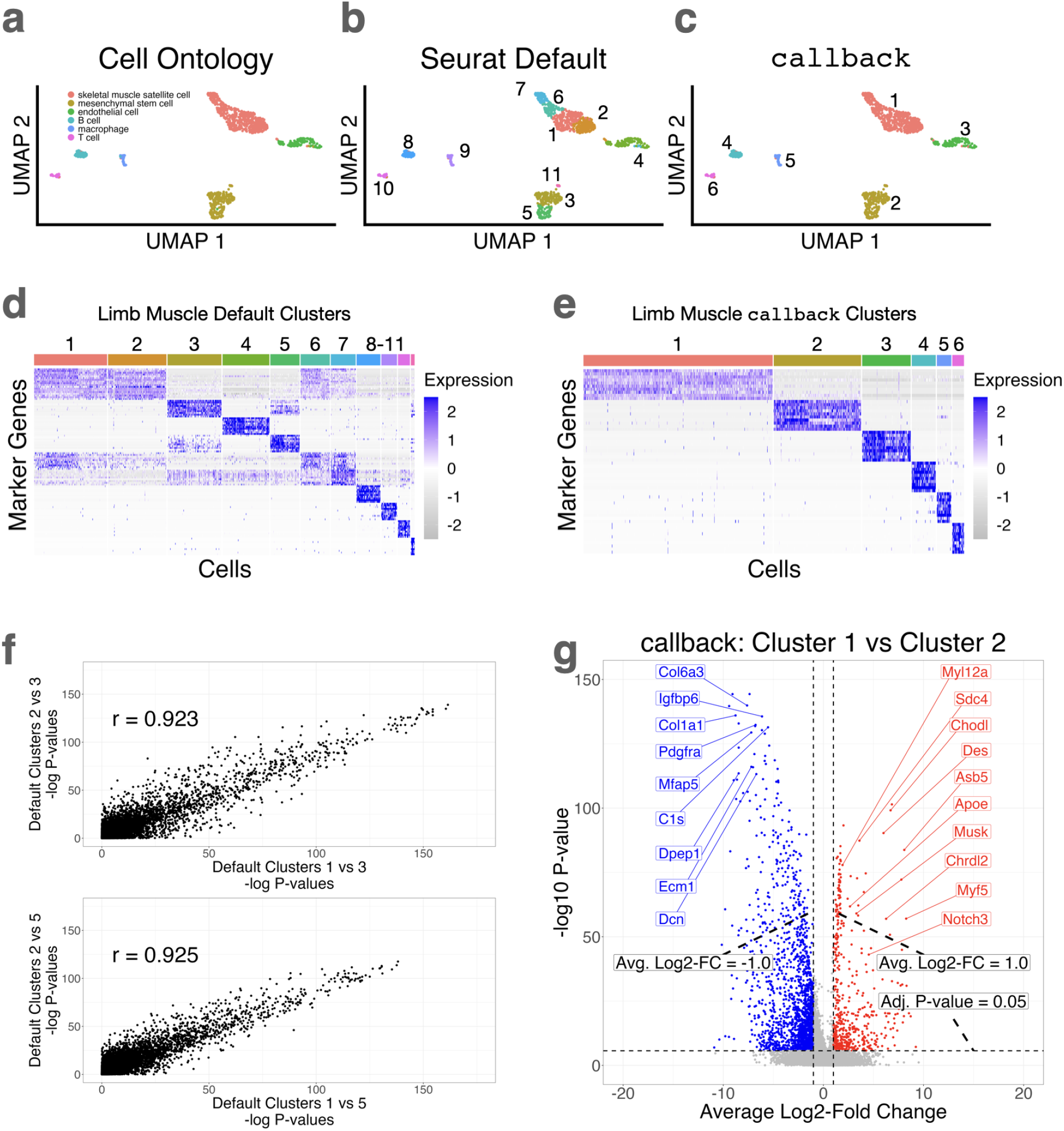
Using callback to avoid over-clustering leads to improved hypothesis generation for downstream analyses. **(a-c)** Uniform manifold approximation and projection (UMAP) plots of **(a)** the manually curated cell ontology class labels, **(b)** inferred clusters using the Louvain algorithm with default parameter settings in `Seurat`, and **(c)** inferred clusters using the Louvain algorithm paired with `callback` for the limb muscle tissue from the Tabula Muris study. **(d)** Heatmap of the top 10 marker genes for each inferred cluster shown in panel **b** with the default Louvain implementation. **(e)** Heatmap of the top 10 marker genes for each inferred cluster shown in panel **c** with the Louvain algorithm paired with `callback`. **(f)** Scatter plots and corresponding Pearson correlation coefficient (*r*) of the log_10_*P* -values for all genes being tested for differential expression between (i) inferred clusters 1 and 2 versus 3 (*r* = 0.923) and (ii) inferred clusters 1 and 2 versus 5 (*r* = 0.925) from panel **(d)** using the default Louvain algorithm in `Seurat`. **(g)** Volcano plot of all genes being tested for differential expression between inferred clusters 1 and 2 from panel **(e)** using the `callback` version of the Louvain algorithm. The genes colored in red and blue are those with a significant *P* -value after Bonferroni correction and with a log_2_-fold change greater than 1 (i.e., up-regulated in cluster 1) or less than -1 (i.e., up-regulated in cluster 2), respectively. The inferred cluster 1 from `callback` corresponds to skeletal muscle satellite cells and cluster 2 corresponds to mesenchymal stem cells. The genes that are labeled are well-known markers of both skeletal muscles (red, up-regulated in cluster 1 relative to cluster 2) and cardiac mesenchymal stem cells (blue, up-regulated in cluster 2 relative to cluster 1).

As a final analysis of computational scalability, we benchmarked the runtime and peak memory use of `callback`, `sc-SHC`, and `CHOIR` on several other publicly available datasets containing 2700, 8444, 30K, and 40K cells (Figs. S26-S27) [17–20]. Each method was run on a machine with 16 cores (Methods). On these datasets, `sc-SHC` was the fastest, closely followed by `callback`, and `CHOIR` was an order of magnitude slower. Additionally, we applied each method using their default settings on subsets of the 68,579 total peripheral blood mononuclear cells (PBMCs) provided by Zheng et al. [1]. These subsets were of sizes 1K, 2K, 5K, 10K, 20K, 30K, 40K, 50K, and 60K cells as well as the full dataset. On these subsets, both `callback` and `sc-SHC` were very similar in speed, while CHOIR was an order of magnitude slower (Fig. S28). In terms of peak memory consumption, `callback` used the least memory while `sc-SHC` showed quadratic memory growth as a function of the number of cells (Fig. S29). In summary, `callback` is as fast (or faster) than alternatives and uses less memory. Notably, `callback` required less than 10 gigabytes (GB) of memory on datasets with nearly 70K cells and was able to cluster those cells in less than 15 minutes (with 16 cores). This demonstrates the ability to analyze large datasets with `callback` on a personal laptop.

The `callback` approach is not without its limitations. First, the algorithm works downward from an upper bound on the number of clusters (often parameterized by *K* in the literature). This strategy could potentially lead to under-clustered results if the starting upper bound is too conservative (i.e., if *K* is too small). To circumvent this limitation, `callback` can be initialized with a large set of clusters; however, this will come with an additional computational cost because more iterations will likely need to be performed until the algorithm converges onto a statistically appropriate number of clusters. Second, the current implementation of `callback` does not account for additional metadata or confounding that might be present in a scRNA-seq dataset. For example, in the presence of batch effects, spurious relationships between cells can be created and `callback` might determine that cells of the same type need to be partitioned into different groups (or vice versa). To that end, incorporating data integration steps, like batch effect correction, into the `callback` software is a relevant direction for future work. One possible extension of the `callback` algorithm would be to run an integration approach (e.g., `Harmony` [21]) on the principal component embeddings of the augmented count matrix to correct for possible confounding before building a KNN graph and performing calibrated clustering.

In conclusion, we have presented `callback`, a novel approach aimed to protect against over-clustering when analyzing single-cell transcriptomic data. Through the analysis of several large-scale datasets, we have shown that `callback` provides state-of-the-art clustering results at a fraction of the runtime and computer memory when compared to other competing algorithms. Importantly, `callback` can be efficiently run on a personal laptop when analyzing tens of thousands of cells. As a disclaimer, cells may exhibit a variety of heterogeneous cell states, continuous axes of variation rather than discrete groups, or other complexities for which `callback`, or any clustering algorithm, is not completely well-suited. Overall, we envision that `callback` will be a useful aid when needing to assign labels to unknown cell types. With both its speed and flexibility, `callback` will save practitioners the hours often spent manually investigating and re-clustering single-cell RNA sequencing datasets.

## Methods

### Overiew of the callback algorithm

Consider a study with single-cell RNA sequencing (scRNA-seq) expression data for *i* = 1, …, *N* cells that each have measurements for *j* = 1, …, *G* genes. Let this dataset be represented by an *N × G* matrix **X** where the column-vector **x**_*j*_ denotes the expression profile for the j-th gene. The `callback` method augments the real expression matrix with knockoff genes which are generated to have no association with any particular cell type [11, 12]. These negative control variables go through the same preprocessing, clustering, and differential expression analyses as the real observed genes in the study; therefore, they are presented with the same opportunity to be identified as marker genes. Since the knockoff genes are essentially noise variables, the distribution of their test statistics represent the impact of post-selective inference (i.e., deviations from the null). As a result, we can correct for these same deviations from the null in the observed test statistics for the real genes which allows us to also calibrate our cluster assignments. This process is also known as implementing a “knockoff filter” (which controls the false discovery rate) when testing for differentially expressed genes between clusters [11, 12]. If there are no detectable differences between the inferred clusters, we assume that over-clustering has occurred and re-cluster with a smaller number of groups.

More specifically, `callback` works by implementing the following steps:

1. For each gene in the study **x**_*j*_, generate a knockoff expression vector 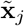. Next, concatenate all of the knockoff genes together and construct a matrix of knockoff variables 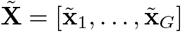 .
2. Combine the real gene expression matrix with the knockoff features into a single object 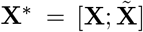 . Then perform the usual preprocessing on the augmented data matrix **X**^*∗*^. In this paper, preprocessing consists of normalizing the expression counts followed by principal component analysis (PCA).
3. Apply a given clustering algorithm (e.g., the Louvain algorithm) to the PCA embeddings of the augmented matrix **X**^*∗*^ (or, alternatively, apply the clustering algorithm to the augmented matrix directly).
4. Conduct differential expression analysis between each *k*-th and *l*-th cluster pair, denoted by *𝒞*_*k*_ and *𝒞*_*l*_, respectively. Obtain *P* -values for all genes (real and knockoff) across each comparison.
5. Let *p*_*j*_(*k*; *l*) represent the *P* -value for the *j*-th real gene when comparing differential expression between clusters *𝒞*_*k*_ and *𝒞*_*l*_. Similarly, let 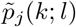 represent the *P* -value for the same comparison but for the corresponding *j*-th knockoff gene. We use these two *P* -values to compute the following knockoff test statistic

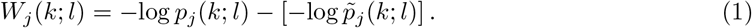

Intuitively, a large, positive value of *W*_*j*_(*k*; *l*) represents evidence that the *j*-th gene is truly different between clusters *𝒞*_*k*_, *𝒞*_*l*_, while a value less than or equal to zero represents strong evidence that there is no difference in the expression of the *j*-th gene between the groups.
6. Next, compute the data-dependent threshold via the following formulation

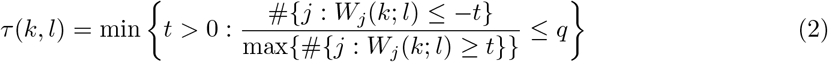

where #*{•}* denotes the cardinality of a set and *q* is a hyperparameter representing the desired false discovery rate (FDR) when testing for differential expression. By default, and for all results presented in this paper, `callback` sets *q* = 0.05. If no such *t >* 0 exists, we set *τ* (*k, l*) = *∞*.

If, for any pair of clusters, *τ* (*k, l*) = *∞*, we return to step #3 and rerun the clustering algorithm with a smaller number of clusters. However, if *τ* (*k, l*) *< ∞* for all pairs of clusters, then we see no evidence of over-clustering and return the inferred cluster assignments to the user.

### Knockoff test statistics

To compute the knockoff test statistics for each cluster *W*_*j*_(*k*; *l*) in Eq. (1), `callback` uses *P* -values *p*_*j*_(*k*; *l*) and 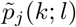 from the Wilcoxon rank sum test as implemented by the `FindMarkers` function in the `Seurat` software package [4] and accelerated by `Presto` [22].

### Differences between callback and ClusterDE

Both `callback` and `ClusterDE` [6] use synthetic null variables and the knockoff filter. The key distinction between these methods is that `ClusterDE` takes given cell clusters and computes knockoff data to calibrate statistical null hypothesis tests between those clusters, while `callback` computes knockoff data on the full dataset first and uses the augmented data matrix as input to the clustering algorithm in order to calibrate the choice of clusters.

### Construction of knockoff genes

To construct knockoff genes that “match” the distribution of expression for the original real genes (but without being associated with any particular cell types), we use a univariate parametric modeling approach which we apply to each individual gene separately. There has been a large body of work focused on choosing the correct distributions for modeling scRNA-seq count data [23–26]. Here, we utilize the zero-inflated Poisson (ZIP) model. Importantly, this parametric generative method creates knockoff gene variables that (i) do not have any association with any particular cell group and (ii) do not retain any covariance structure with the original real genes. The ZIP model mixes two generative processes—the first generates zeros and the second is governed by a Poisson distribution that generates counts (some of which may also be zero) [27]. For a random variable *X ∼* ZIP(*π*_0_, *λ*), we have the following mixture

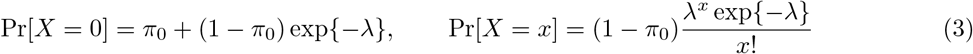

where *x ∈* ℕ ^+^ is any non-negative integer value, *λ* is the expected count from the Poisson distribution (i.e., the rate parameter), and *π*_0_ is the proportion of extra zeroes arising in addition to those from the underlying Poisson distribution. The maximum likelihood estimators for the ZIP model, given the expression of the *j*-th gene, take the following form

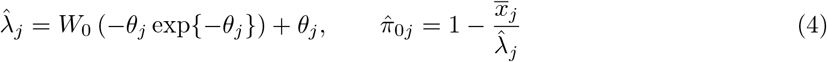

where 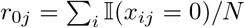 denotes the proportion of observed zeroes for the *j*-th gene across all cells (with 𝕀 (*•*) being an indicator function), 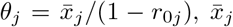 is the sample average expression for the *j*-th gene of interest, and *W*_0_ is the principal branch of the Lambert *W* function (i.e., *W*_0_(*a*) = *b* implies *b* exp*{b}* = *a*). For each *j*-th real gene **x**_*j*_, we fit the maximum likelihood estimators 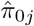 and 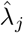 and then sample the synthetic expression for the corresponding knockoff gene as 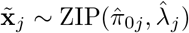.

### Parameters for the callback algorithm

The default starting resolution parameter for the Louvain and Leiden algorithms within `callback` is *γ* = 0.8, the same as the default in the `FindClusters` function in `Seurat`. Since `callback` works by iteratively reducing the starting number of clusters, if the starting parameter is too low (i.e., if you start with correctly calibrated clusters or under-cluster) there is no opportunity for `callback` to iteratively reduce the number of clusters. There is a warning produced by `callback` software when this occurs and users can re-run `callback` with a new parameter to begin with a larger number of clusters.

### Simulation study

We simulated scRNA-seq data using the `splatter R` package [28] which implements a gamma-Poisson model to create a count matrix for cells. In Fig. 1, the one-group dataset was simulated with 1000 genes and 1000 cells; while the three-group dataset was simulated to have 1000 genes and 4000 cells with the three groups being separated in proportions of 0.6, 0.2, and 0.2, respectively. Differential gene expression between the groups was controlled using the `de.prob` parameter with a value 0f 0.05.

### Preprocessing and data availability

Below we briefly describe all of the datasets used in this work. All datasets outside of the Tabula Muris were used exclusively to test the scalability of `callback` and competing methods; therefore, clustering performance was not recorded. All preprocessing steps were done using the `Seurat` software package. For each of these datasets, the count matrices were log-normalized using the `NormalizeData` function with the default parameters. Here, we set the `scale.factor` = 10000. The number of variable genes was set to 1000 for all analyses. These were determined by using the `vst` selection method implemented by the `FindVariableFeatures` function. All data were centered and scaled using the `ScaleData` function with default parameters, principle components were computed with the `RunPCA` using the variable genes as input, and the nearest neighbor graphs were computed using the first 10 principal components within the `FindNeighbors` function. Each evaluated method (`callback`, `sc-SHC`, and `CHOIR`) was provided with the top 1000 highly variable genes and the first 10 principal component embeddings. The implementations of the Louvain clustering algorithms analyzed the nearest neighbor graphs with resolution values set to *γ* = 0.8.

### Tabula Muris

To compare the clustering performance of `callback` against competing methods, we utilized the 20 organs from the Tabula Muris dataset [13]. This dataset contains 53,760 total cells with human-curated cell type labels for each organ. After following the quality control steps outlined in the original study (i.e., filtering to exclude cells with less than 500 total genes detected and to exclude cells with less than 50,000 total reads) and additionally removing cells without a manually curated cell type label, we were left with a total of 45,423 cells for the analysis. The individual scRNA-seq expression datasets for each tissue can be found on figshare: https://figshare.com/articles/dataset/Single-cell_RNA-seq_data_from_Smart-seq2_sequencing_of_FACS_sorted_cells/5715040.

### PBMC 3K, Bone Marrow 30K, and Bone Marrow 40K

To assess the runtime and peak memory usage of `callback` and other competing approaches, we utilized multiple datasets available through the `SeuratData` R package found here: https://github.com/satijalab/seurat-data. In particular, we downloaded data under the `pbmc3k`, `bmcite`, and `hcabm40k` variable names. For each of these datasets, `callback` was run with a larger starting resolution parameter of *γ* = 1.5 to ensure that more than one iteration took place.

### PBMC 68K

We took scRNA-seq data from fluorescence-activated cell sorted (FACS) populations of peripheral blood mononuclear cells (PBMCs) provided by Zheng et al. [1] and concatenated each population into one dataset. This dataset contains 68,579 cells with ten different labels corresponding to each purified population that was sorted. The dataset can be found on the 10X Genomics website and the URL can be found on this GitHub page: https://github.com/10XGenomics/single-cell-3prime-paper/blob/master/pbmc68k_analysis/README.md. It can also be directly downloaded here: https://cf.10xgenomics.com/samples/cell/pbmc68k_rds/pbmc68k_data.rds.

### Liver 8K

This dataset contains 8,444 cells provided by MacParland et al. [18]. It can be loaded using the `HumanLiver R` package available here: https://github.com/BaderLab/HumanLiver. For this dataset, `callback` was run with a larger starting resolution parameter of *γ* = 1.5 to ensure that more than one iteration took place.

## Code availability

All code is available under the open-source MIT license at https://github.com/lcrawlab/callback with documentation at https://lcrawlab.github.io/callback. The scripts used to analyze the data and to reproduce the figures from this paper are available at https://github.com/lcrawlab/callbackreproducibility. The fully rendered results can also be viewed at https://lcrawlab.github.io/callbackreproducibility.

## Acknowledgements

We thank members of the Crawford, Shalek, Raghavan, and Winter Labs for insightful comments on earlier versions of this manuscript. This research was conducted using computational resources and services provided by the Center for Computation and Visualization at Brown University. This research was also supported in part by a David & Lucile Packard Fellowship for Science and Engineering awarded to LC. Any opinions, findings, and conclusions or recommendations expressed in this material are those of the author(s) and do not necessarily reflect the views of any of the funders.

## Author contributions

AD and LC conceived the study and developed the methods. AD developed the algorithm, software, and led the analyses. AD, MLR, and AWN conducted secondary analyses. SR, PSW, APA, and LC co-supervised the project. AKS and LC provided resources. AD and LC wrote the initial draft. All authors interpreted the results, and revised the manuscript.

## Competing interests

SR holds equity in Amgen. SR and PSW receive research funding from Microsoft. AKS reports compensation for consulting and/or scientific advisory board membership from Honeycomb Biotechnologies, Cellarity, Ochre Bio, Relation Therapeutics, Fog Pharma, Bio-Rad Laboratories, IntrECate Biotherapeutics, Passkey Therapeutics and Dahlia Biosciences unrelated to this work. All other authors have declared that no competing interests exist.

## Notes

https://github.com/lcrawlab/callback

https://lcrawlab.github.io/callback/

https://github.com/lcrawlab/callbackreproducibility

https://lcrawlab.github.io/callbackreproducibility/

