## Supplemental Material for "A knockoff calibration method to avoid over-clustering in single-cell RNA-sequencing"

### Supplementary Material for "A knockoff calibration method to avoid over-clustering in single-cell RNA-sequencing"

#### Cluster Evaluation Metrics Vignette

In this Supplementary Note, we detail the metrics used to evaluate the quality of clustering assignments against ground truth, which are referred to as extrinsic clustering metrics. For the analyses conducted in the main text, we use the manually curated cell type annotations from the original studies as a proxy for ground truth labels. All of the metrics described here (except for the adjusted Rand index) vary between 0 (poor agreement between inferred cluster assignments and ground truth) and 1 (good agreement between inferred cluster assignments and ground truth). The adjusted Rand index (ARI) ranges between [-1/2, 1] where 1 represents perfect agreement between label sets, 0 represents random agreement, and negative values represent worse than random agreement [1]. There are two broad classes of clustering evaluation metrics utilizing ground truth labels: (1) confusion matrix-based metrics which are based on notions of true positives, true negatives, false positives, and false negatives, and (2) entropy-based metrics which measure the uncertainty in inferred clustering assignments given a set of reference labels (and vice versa).

##### Confusion Matrix-Based Cluster Evaluation Metrics

The typical confusion matrix is defined as follows for a classification problem with 2 classes.

| | | True Label | | |
|---|---|---|---|---|
| | | Class 1 | Class 2 | Total |
| Inferred Label | Class 1 | TP | FP | $a_1$ |
| | Class 2 | FN | TN | $a_2$ |
| | Total | $b_1$ | $b_2$ | $n$ |

where TP is the number of true positives, FP is the number of false positives, TN is the number of true negatives, and FN is the number of false negatives. This can be similarly defined for a classification problem with an arbitrary number of classes. Since a clustering result may have a different number of inferred clusters than the number of true labels and since it is not immediately obvious how to know which true label an inferred cluster might correspond to, we instead define these metrics over pairs of points in the following way:

- True positives (TP) is the number of cell pairs that have the **same** true labels and are assigned the **same** inferred labels after clustering.
- True negatives (TN) is the number of cell pairs that have **different** true labels and also have **different** inferred labels after clustering.

- False positives (FP) is the number cell pairs that have **different** true labels but are assigned the **same** inferred labels after clustering.
- False negatives (FN) is the number of cell pairs that have the **same** true labels but are assigned **different** inferred labels after clustering.

We describe a series of confusion matrix-based cluster evaluation metrics below using these concepts. To simplify notation, we will let $n_{ij}$, $a_i$, and $b_j$ be values obtained from a contingency table, and $n = \sum_{ij} n_{ij}$ for indices $i$ and $j \in \{1, 2\}$. To be specific, $\text{TP} = n_{11}$ , $\text{FP} = n_{12}$, $\text{FN} = n_{21}$, and $\text{TN} = n_{22}$, respectively. An illustration of these metrics can be found in Fig. S1.

##### Adjusted Rand Index (ARI)

This adjusted Rand index (ARI) captures the similarity between labels inferred by a clustering algorithm and the reference labels. It is based on the Rand index (RI) which is computed as the following

$$\text{RI} = \frac{\text{TP} + \text{TN}}{\text{TP} + \text{FP} + \text{FN} + \text{TN}} = \frac{b_1}{n}. \tag{1}$$

The ARI corrects for the RI measurement's sensitivity to chance via permutation [2] where

$$\text{ARI} = \frac{\text{RI} - \mathbb{E}[\text{RI}]}{1 - \mathbb{E}[\text{RI}]} = \frac{\sum_{ij} \binom{n_{ij}}{2} - \left[\sum_i \binom{a_i}{2} \sum_j \binom{b_j}{2}\right] \binom{n}{2}}{1/2 \left[\sum_i \binom{a_i}{2} + \sum_j \binom{b_j}{2}\right] - \left[\sum_i \binom{a_i}{2} \sum_j \binom{b_j}{2}\right] \binom{n}{2}}. \tag{2}$$

Here, $\mathbb{E}[\text{RI}]$ denotes the expectation of the Rand index.

##### Jaccard Similarity Index

For high-dimensional single-cell studies with a large number of both cells and cell types, the number of true negatives can be quite large. The Jaccard similarity index captures the amount of overlap between two finite sets. Overall, the Jaccard index can be a useful alternative to the ARI because it ignores the contribution of the true negatives. Formally, it is defined as the following

$$J = \frac{\text{TP}}{\text{TP} + \text{FP} + \text{FN}} = \frac{n_{11}}{a_1 + n_{21}} \tag{3}$$

which ranges between [0, 1] where 1 denotes complete overlap between the inferred and true cluster labels, and 0 signifies no overlap.

##### Folkes-Mallows Index (FMI)

The Folkes-Mallows Index is the geometric mean of the positive predictive value (PPV) and the true positive rate (TPR) [3]. More concretely, these values are computed via the following

$$\text{PPV} = \frac{\text{TP}}{\text{TP} + \text{FP}} = \frac{n_{11}}{a_1} \qquad \text{TPR} = \frac{\text{TP}}{\text{TP} + \text{FN}} = \frac{n_{11}}{b_1} \tag{4}$$

A higher value for the Fowlkes–Mallows index indicates a greater similarity between the inferred clusters and the true labels. This is computed by

$$\text{FMI} = \sqrt{\text{PPV} \times \text{PPR}}. \tag{5}$$

The FMI is also on the unit scale ranging from [0, 1] where 0 corresponds to the worst inferred clusters such that both inferred and true groups are completely unrelated, and 1 corresponds to the scenario in which both groupings perfectly agree.

### Entropy-Based Cluster Evaluation Metrics

Entropy-based clustering metrics aim to quantify the amount of information shared between the inferred label distribution and the reference label distribution. We describe a series of these metrics below.

#### Completeness

Completeness describes the uncertainty in the inferred clustering assignment conditioned on the true labels where 1 is a "good" value for satisfying completeness. In other words, it quantifies how often cells of the same type are also in the same cluster. Formally,

$$C = 1 - \frac{\mathbb{H}(\mathcal{A} \,|\, \mathcal{B})}{\mathbb{H}(\mathcal{A})} \tag{6}$$

where $\mathcal{B}$ represents the set of true labels, $\mathcal{A}$ represents the inferred clustering assignments, and $\mathbb{H}(\cdot)$ represents the Shannon entropy of a given label distribution [4]. When all cells from true group $B$ belong to the same inferred cluster, the completeness metric equals 1. For the degenerate case where $\mathbb{H}(\mathcal{A}, \mathcal{B}) = 0$, we define $C = 0$.

#### Homogeneity

Homogeneity describes the uncertainty in the true label conditioned on the cluster assignment where 1 is a "good" value for satisfying homogeneity. In other words, it quantifies the amount that cells within a cluster are also from the same cell type. Formally, homogeneity is defined

$$H = 1 - \frac{\mathbb{H}(\mathcal{B} \,|\, \mathcal{A})}{\mathbb{H}(\mathcal{B})} \tag{7}$$

where, again, $\mathcal{B}$ represents the set of true labels, $\mathcal{A}$ represents the inferred clustering assignments, and $\mathbb{H}(\cdot)$ represents the Shannon entropy of a given label distribution [4]. When all cells from all of the clusters contain only cells that are from the same true group, the homogeneity metric equals 1. For the degenerate case where $\mathbb{H}(\mathcal{A}, \mathcal{B}) = 0$, we define $H = 0$.

#### *V*-Measure Balances Between Completeness and Homogeneity

A key observation when using completeness and homogeneity to evaluate clustering algorithms is that they tend to penalize over- and under-clustering, respectively.

- Consider a simple example of over-clustering where there is a group of cells that is split in half into two clusters. Then the uncertainty in the inferred clustering assignments $\mathcal{A}$, given the true labels $\mathcal{B}$, is high because half of the group is in each cluster. Formally, this means that the Shannon entropy $\mathbb{H}(\mathcal{A} \,|\, \mathcal{B})$ is high and so the completeness will be low.
- On the other hand, consider a simple example of under-clustering where two different cell groups are assigned to the same cluster. Then the uncertainty in the true group labels $\mathcal{B}$, given the inferred clustering assignments $\mathcal{A}$, is high because the cells in the cluster could come from either of the two groups. Formally, the entropy $\mathbb{H}(\mathcal{B} \,|\, \mathcal{A})$ is high and so homogeneity will be low.

The $V$-measure is defined as the weighted harmonic mean of completeness and homogeneity where

$$V(\beta) = \frac{(1+\beta)HC}{\beta H + C} \tag{8}$$

where $\beta$ dictates the contribution of completeness versus homogeneity [4]. In this paper, we set $\beta = 1$ such that the two are weighted equally and the $V$-measure is simply

$$V(1) = \frac{2HC}{H + C} \tag{9}$$

which is the simple harmonic mean of completeness and homogeneity. The $V$-measure ranges between [0, 1] where 1 represents total information sharing between label sets and 0 represents no information sharing between label sets. It requires that both homogeneity and completeness are maximized and will be zero if a clustering algorithm completely fails to satisfy either of the two properties.

#### Simulation Study: Behavior of Cluster Evaluation Metrics

In order to quantitatively demonstrate the behavior of both the confusion matrix-based and entropy-based clustering metrics, we consider a synthetic dataset with three groups of cells (see Fig. S1). When an algorithm under-clusters, it will merge two of the clusters; while, when an algorithm over-clusters, it will infer a fourth group made up of cells from two of the true groups. The table below shows the behavior of different clustering evaluation metrics in the cases when an algorithm finds the true groups, under-clusters, and over-clusters.

| [2pt] Metric | True Groups | Under-Clustered | Over-Clustered |
|---|---|---|---|
| ARI | 1.0 | 0.55 | 0.83 |
| Jaccard index | 1.0 | 0.58 | 0.78 |
| FMI | 1.0 | 0.76 | 0.88 |
| Homogeneity | 1.0 | 0.54 | 0.94 |
| Completeness | 1.0 | 0.92 | 0.77 |
| $V$-measure | 1.0 | 0.68 | 0.85 |

Notice that ARI, Jaccard index, and homogeneity qualitatively perform similarly by penalizing under-clustering more than over-clustering. In contrast, completeness penalizes over-clustering and rewards under-clustering. FMI and $V$-measure balance between both metrics. For this reason, in the main text, we focus on ARI due to its popularity in the literature [5] and $V$-measure because of its ability to balance the impact of over-clustering and under-clustering when evaluating `callback`, `sc-SHC`, and `CHOIR`.

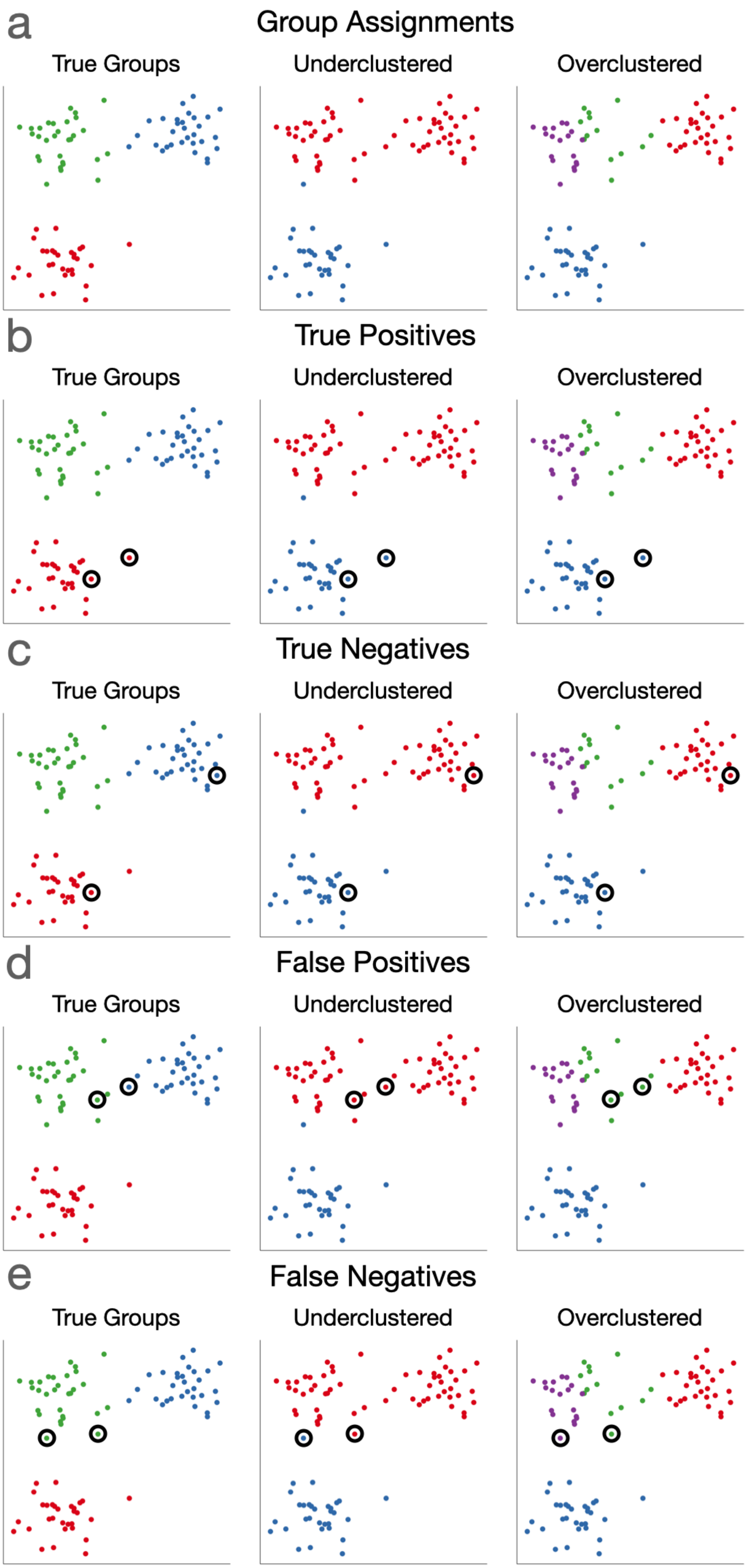


**Figure S1.** **(Continued on the following page).**

**Figure S1. Demonstration of cases of over- and under-clustering in single-cell analyses.** Here, we generate synthetic data from a Gaussian mixture model. Data are created such that there are three true groups. Panel **(a)** shows a depiction of the true cluster labels and the inferred cluster assignments when an algorithm under- and over-clusters, respectively. For comparison, we also show an example of definitions for **(b)** true positives, **(c)** true negatives, **(d)** false positives, and **(e)** false negatives used by confusion matrix-based metrics.

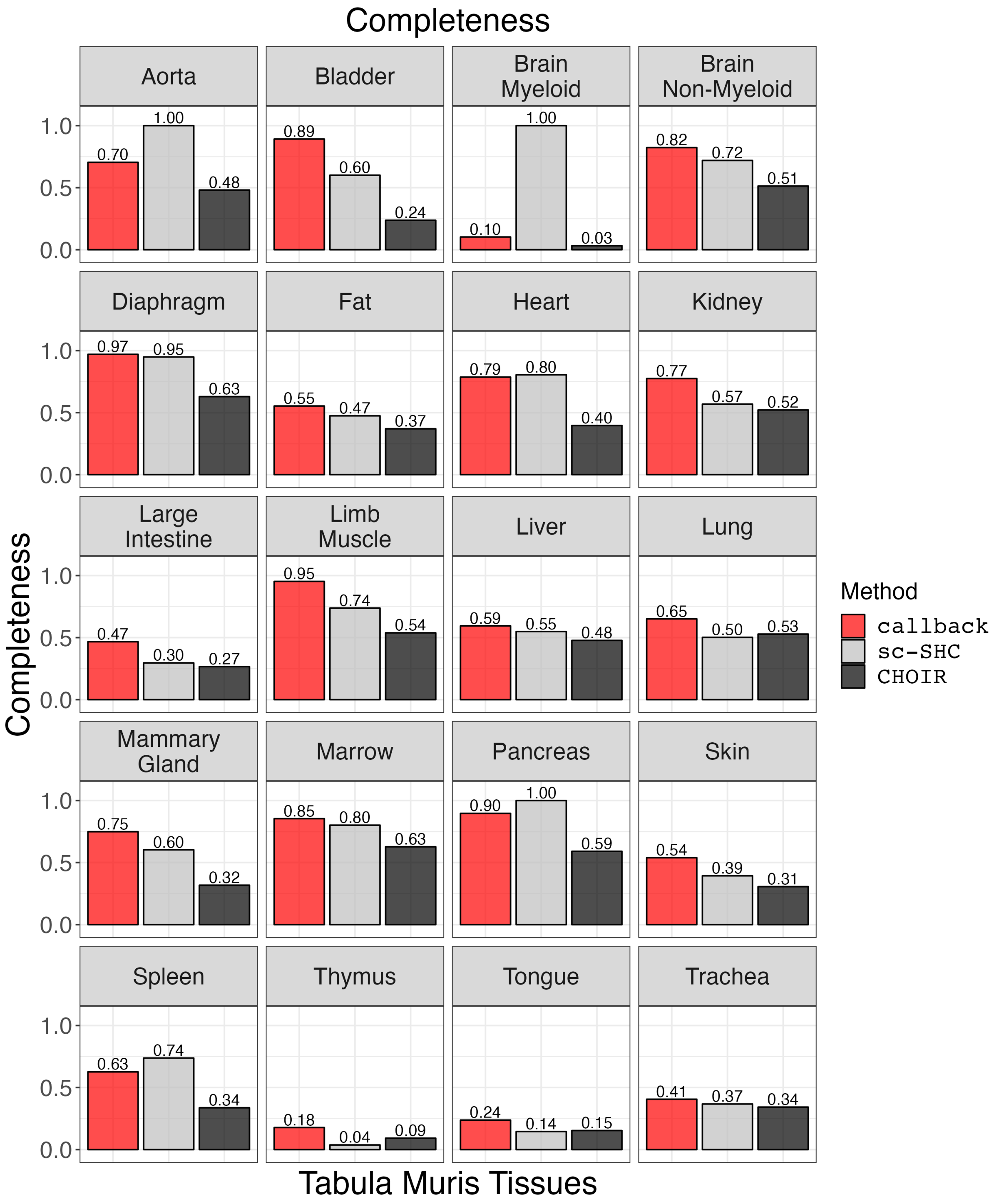


**Figure S2. Performance comparison of `callback`, `sc-SHC`, and `CHOIR` using completeness for each tissue in the Tabula Muris dataset.**

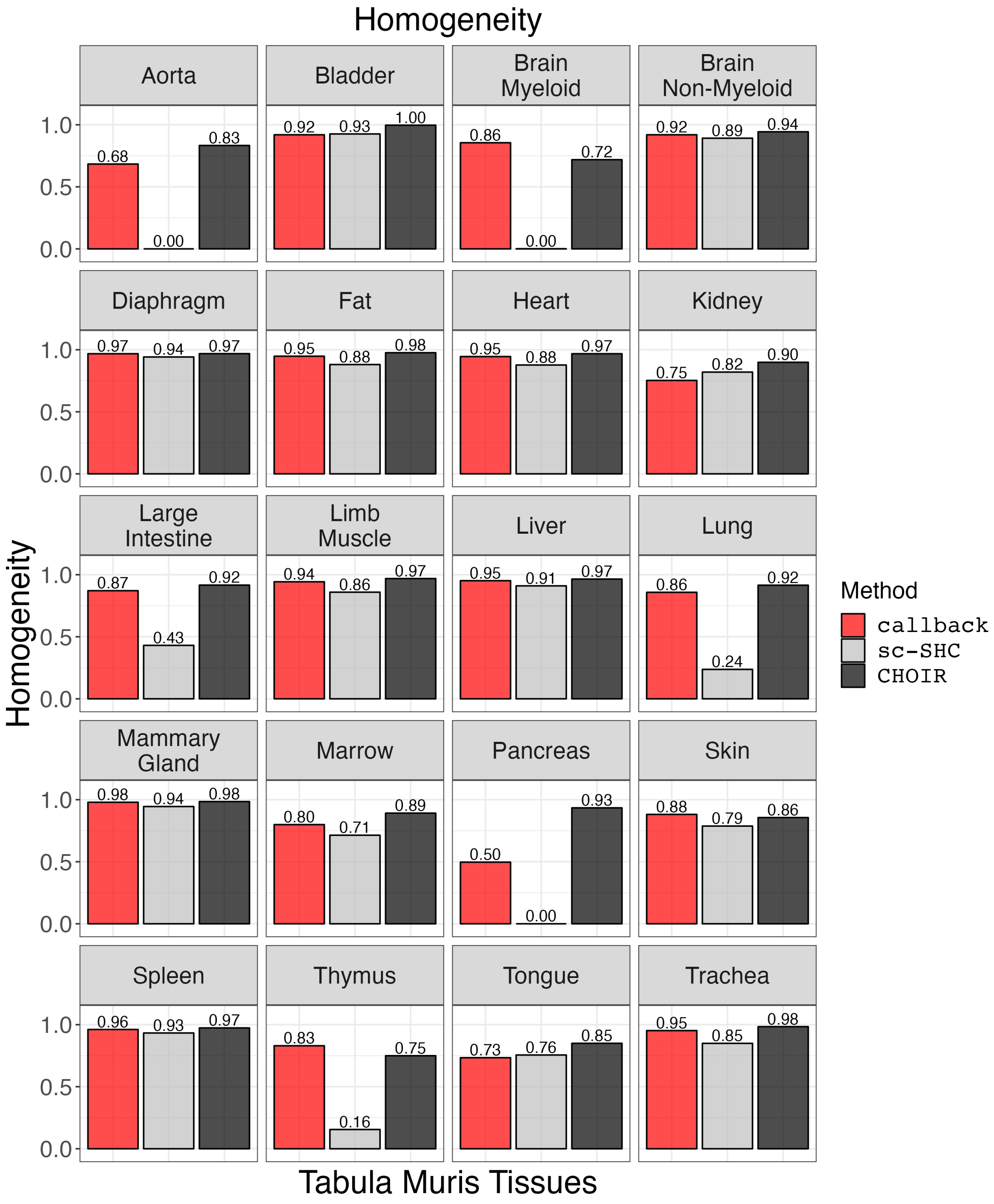


**Figure S3. Performance comparison of `callback`, `sc-SHC`, and `CHOIR` using homogeneity for each tissue in the Tabula Muris dataset.**

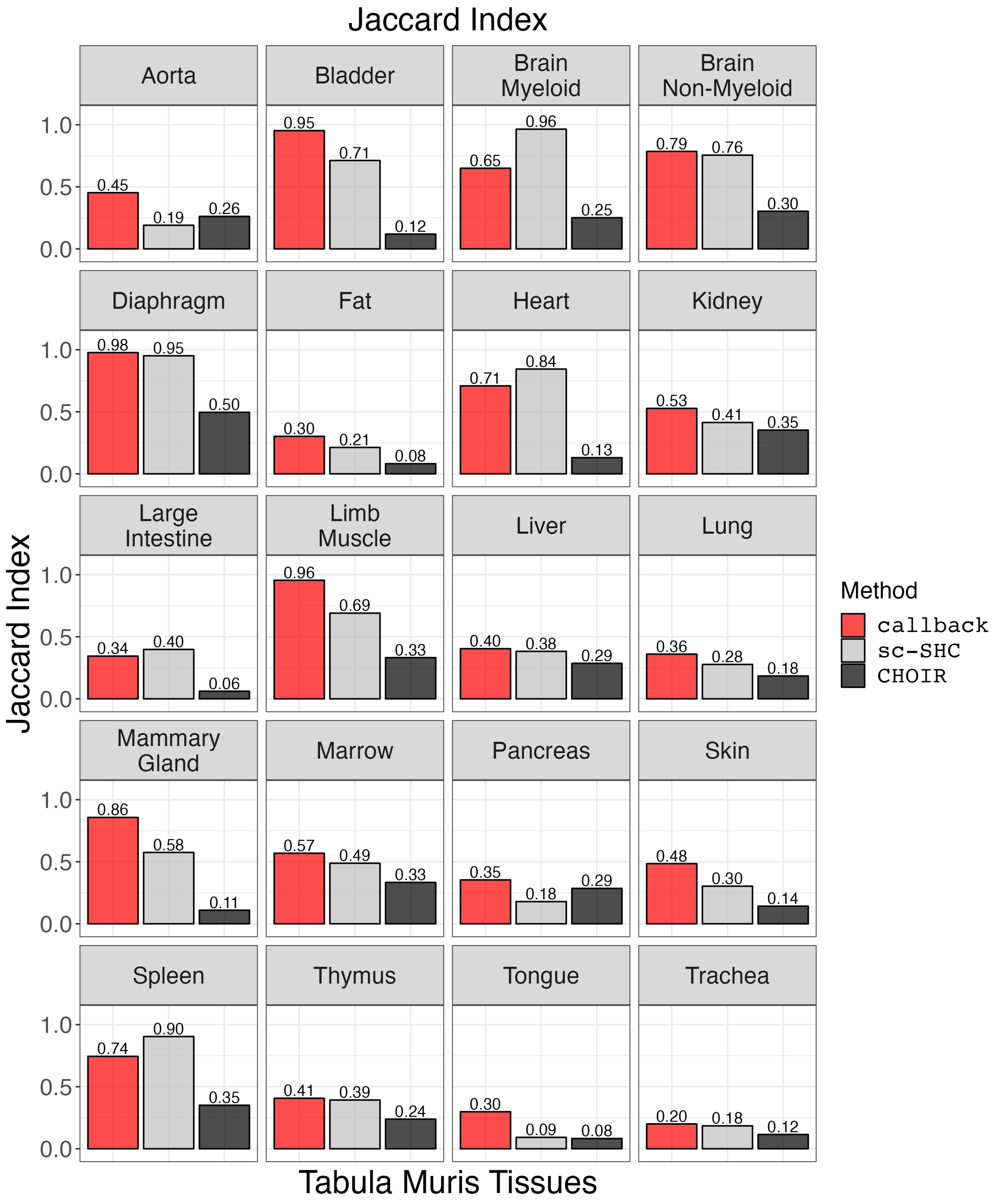


**Figure S4. Performance comparison of `callback`, `sc-SHC`, and `CHOIR` using the Jaccard Index for each tissue in the Tabula Muris dataset.**

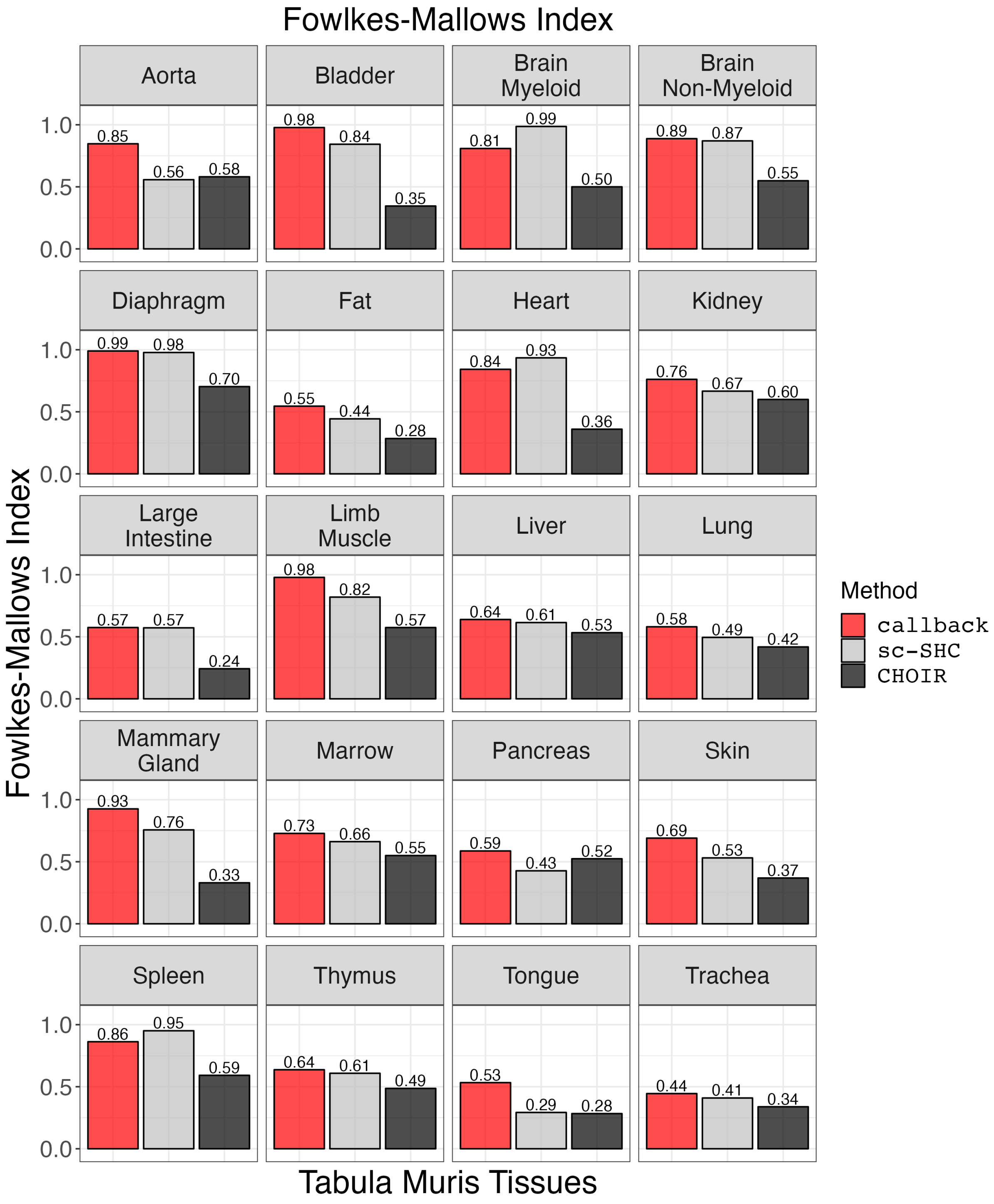


**Figure S5. Performance comparison of `callback`, `sc-SHC`, and `CHOIR` using the Fowlkes-Mallows Index for each tissue in the Tabula Muris dataset.**

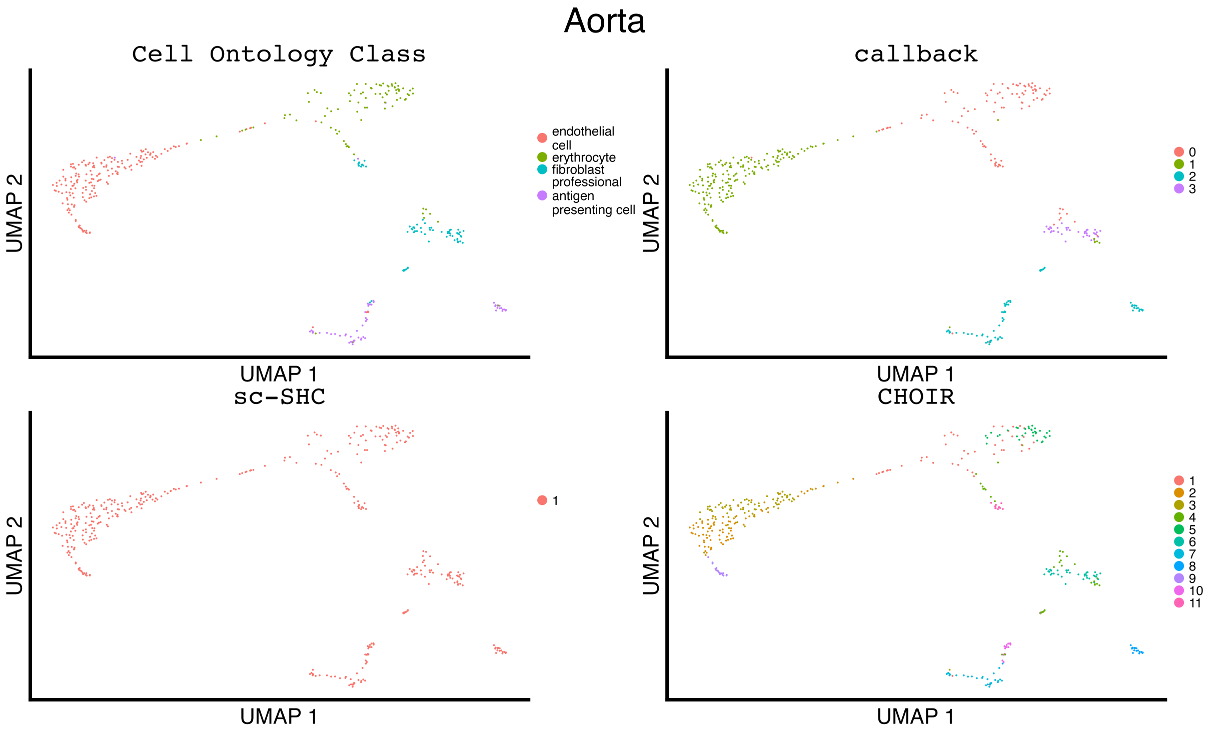


**Figure S6. Uniform manifold approximation and projection (UMAP) plots illustrating the manually curated cell ontology class labels compared to the inferred clustering results for `callback`, `sc-SHC`, and `CHOIR` when analyzing the aorta tissue from the Tabula Muris study.**

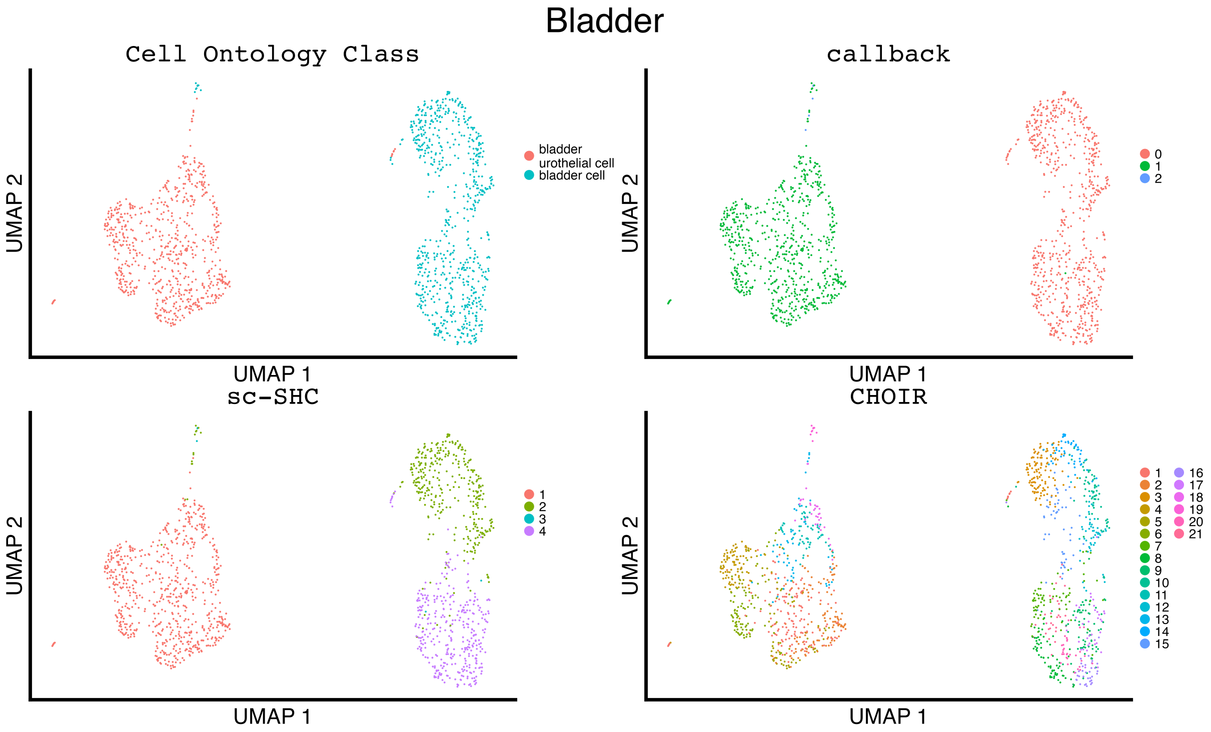


**Figure S7. Uniform manifold approximation and projection (UMAP) plots illustrating the manually curated cell ontology class labels compared to the inferred clustering results for `callback`, `sc-SHC`, and `CHOIR` when analyzing the bladder tissue from the Tabula Muris study.**

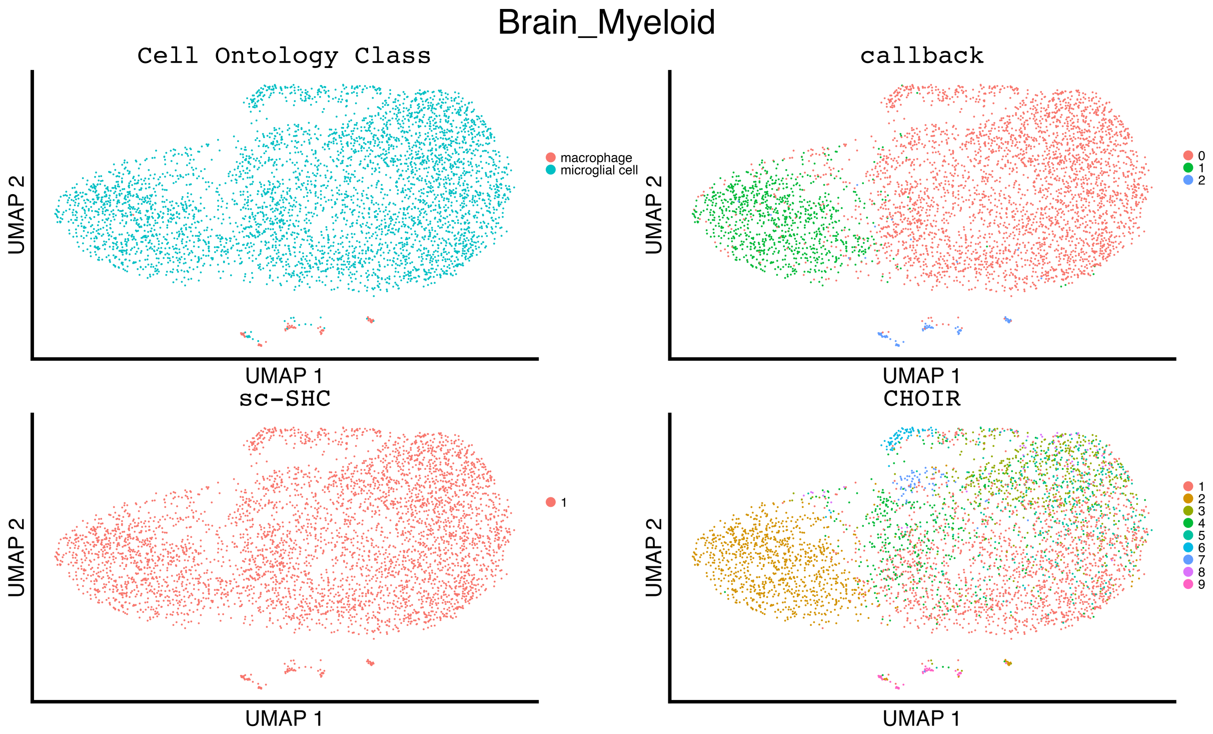


**Figure S8. Uniform manifold approximation and projection (UMAP) plots illustrating the manually curated cell ontology class labels compared to the inferred clustering results for `callback`, `sc-SHC`, and `CHOIR` when analyzing the Brain myeloid tissue from the Tabula Muris study.**

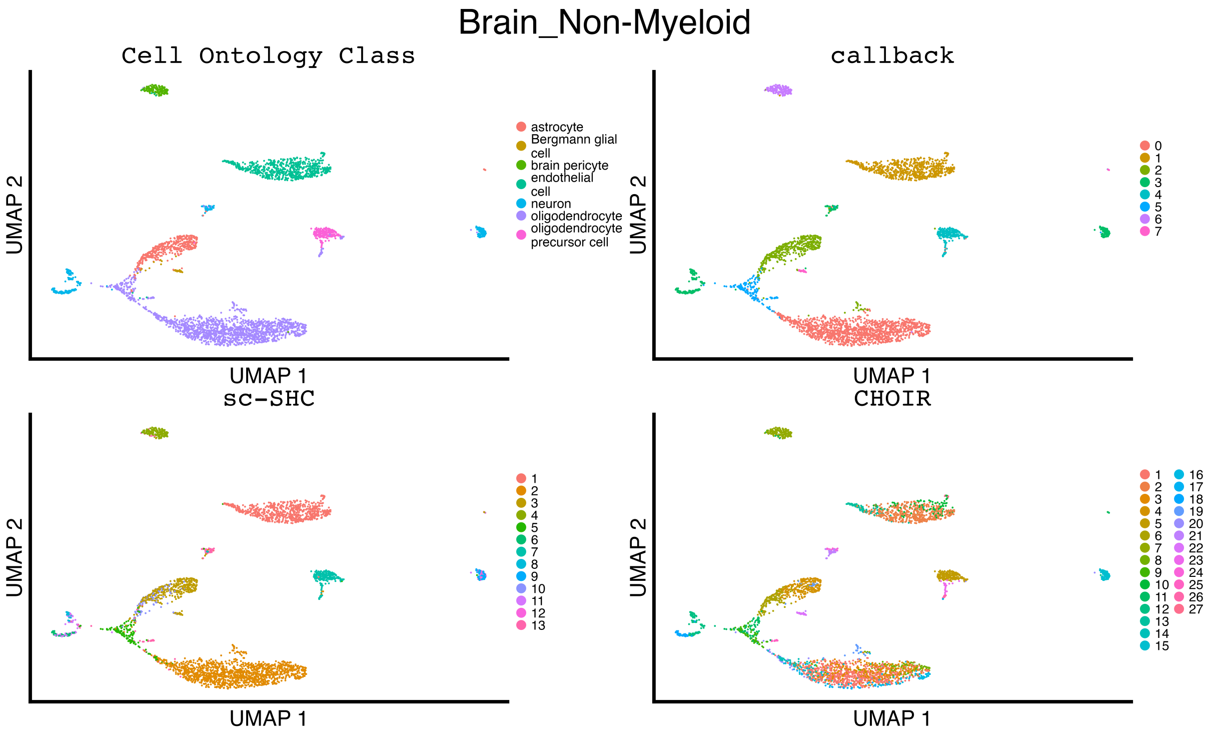


**Figure S9. Uniform manifold approximation and projection (UMAP) plots illustrating the manually curated cell ontology class labels compared to the inferred clustering results for `callback`, `sc-SHC`, and `CHOIR` when analyzing the Brain non-myeloid tissue from the Tabula Muris study.**

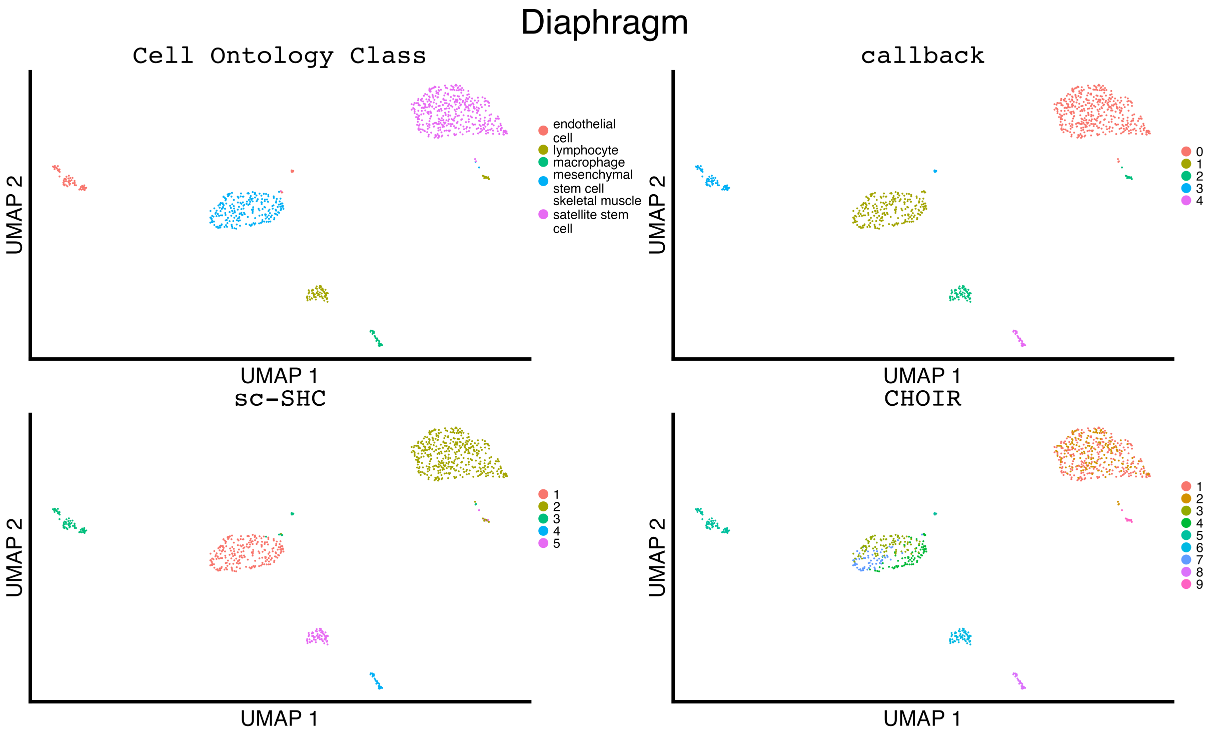


**Figure S10. Uniform manifold approximation and projection (UMAP) plots illustrating the manually curated cell ontology class labels compared to the inferred clustering results for** `callback`**,** `sc-SHC`**, and** `CHOIR` **when analyzing the diaphragm tissue from the Tabula Muris study.**

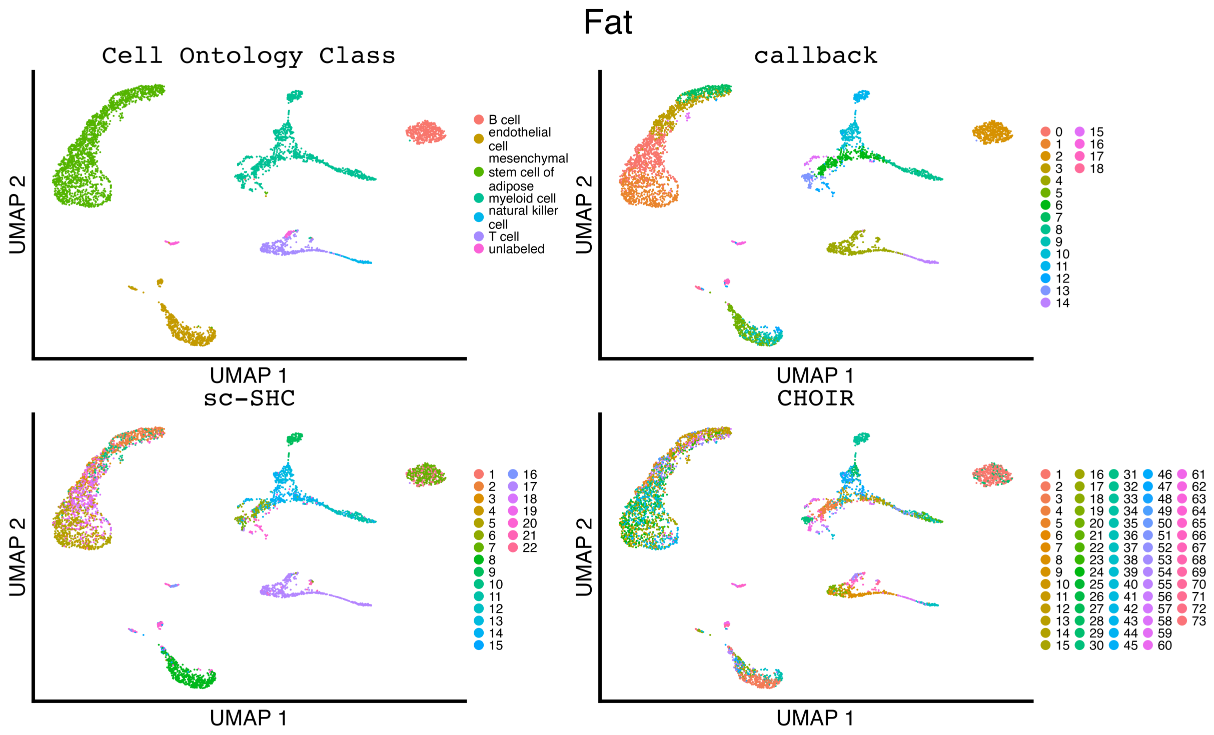


**Figure S11. Uniform manifold approximation and projection (UMAP) plots illustrating the manually curated cell ontology class labels compared to the inferred clustering results for `callback`, `sc-SHC`, and `CHOIR` when analyzing the fat tissue from the Tabula Muris study.**

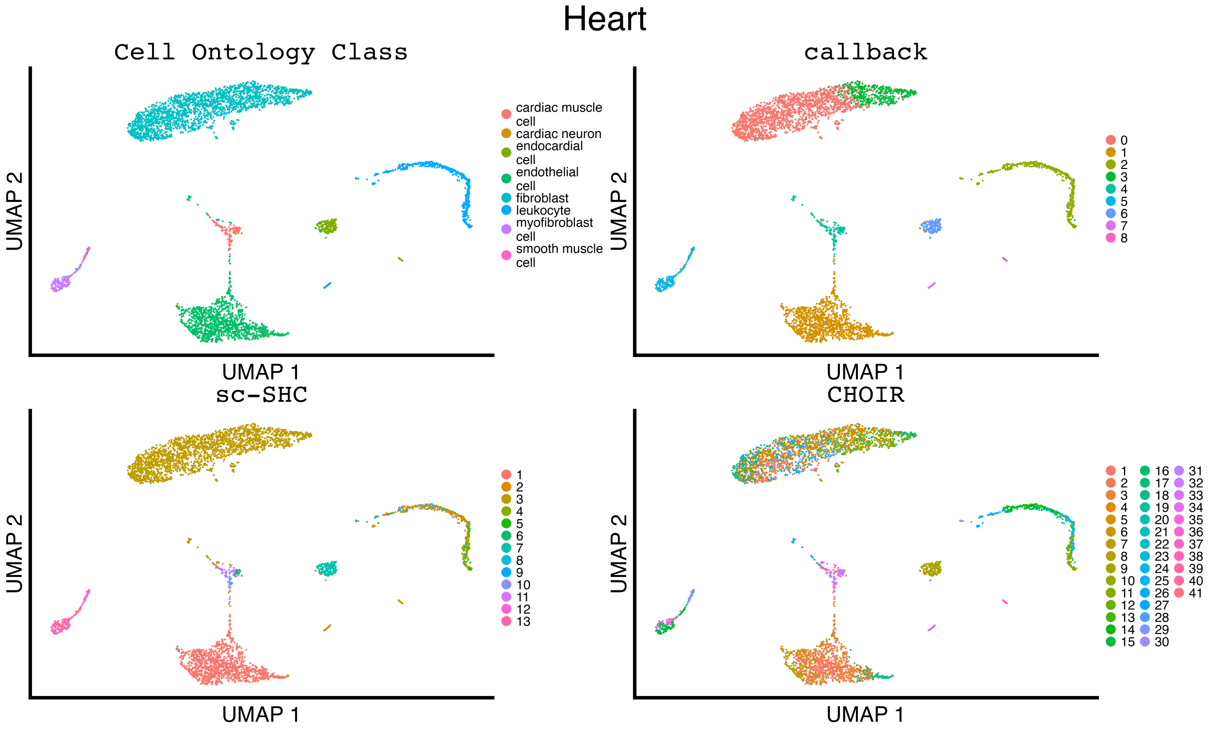


**Figure S12. Uniform manifold approximation and projection (UMAP) plots illustrating the manually curated cell ontology class labels compared to the inferred clustering results for `callback`, `sc-SHC`, and `CHOIR` when analyzing the heart tissue from the Tabula Muris study.**

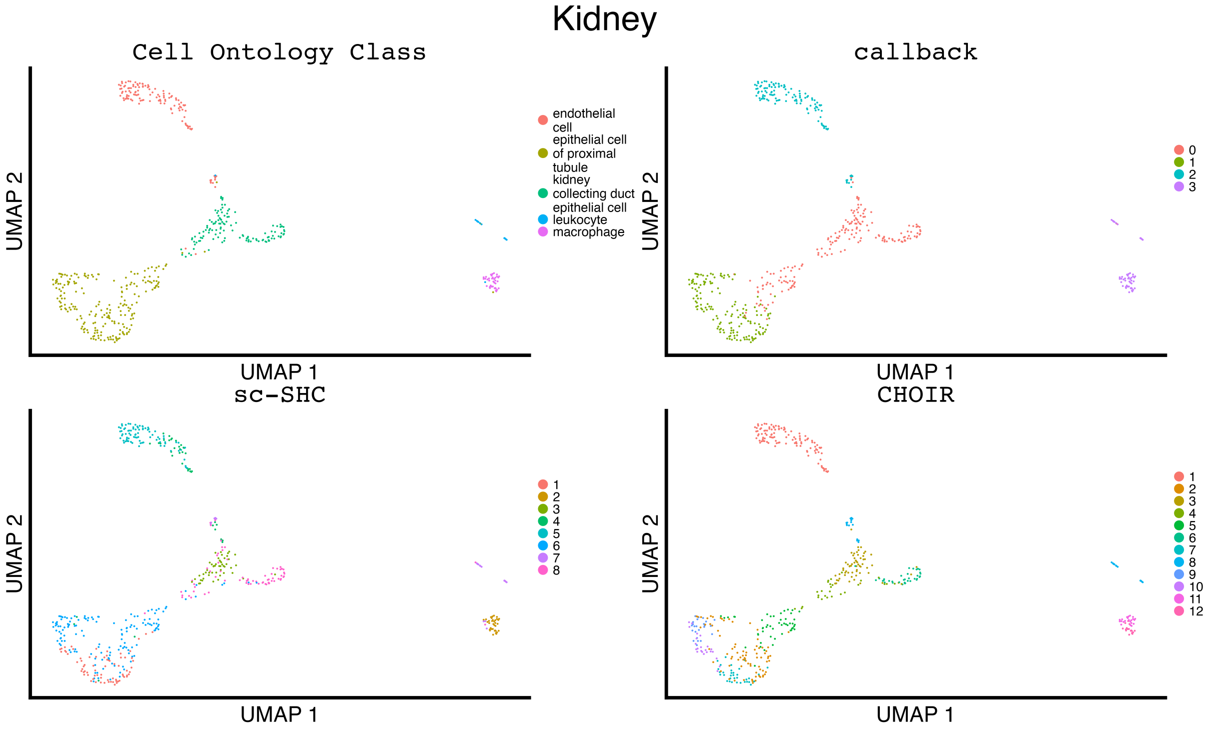


**Figure S13. Uniform manifold approximation and projection (UMAP) plots illustrating the manually curated cell ontology class labels compared to the inferred clustering results for `callback`, `sc-SHC`, and `CHOIR` when analyzing the kidney tissue from the Tabula Muris study.**

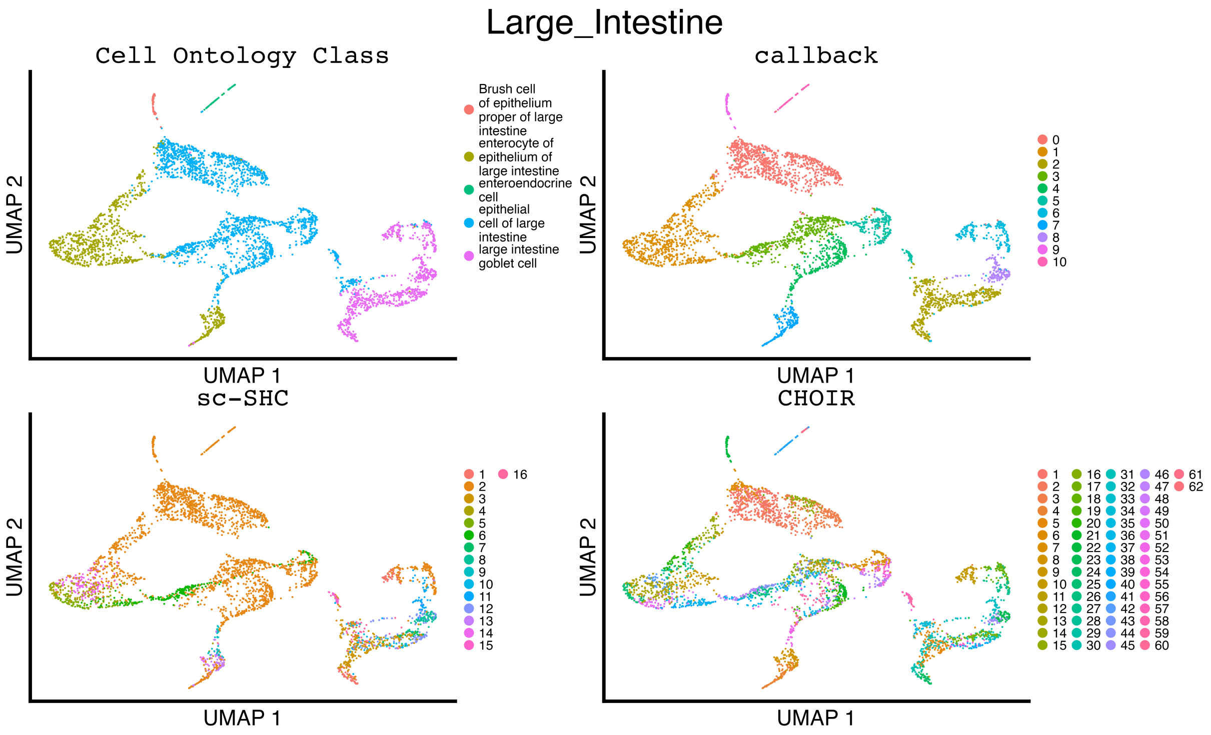


**Figure S14. Uniform manifold approximation and projection (UMAP) plots illustrating the manually curated cell ontology class labels compared to the inferred clustering results for `callback`, `sc-SHC`, and `CHOIR` when analyzing the large intestine tissue from the Tabula Muris study.**

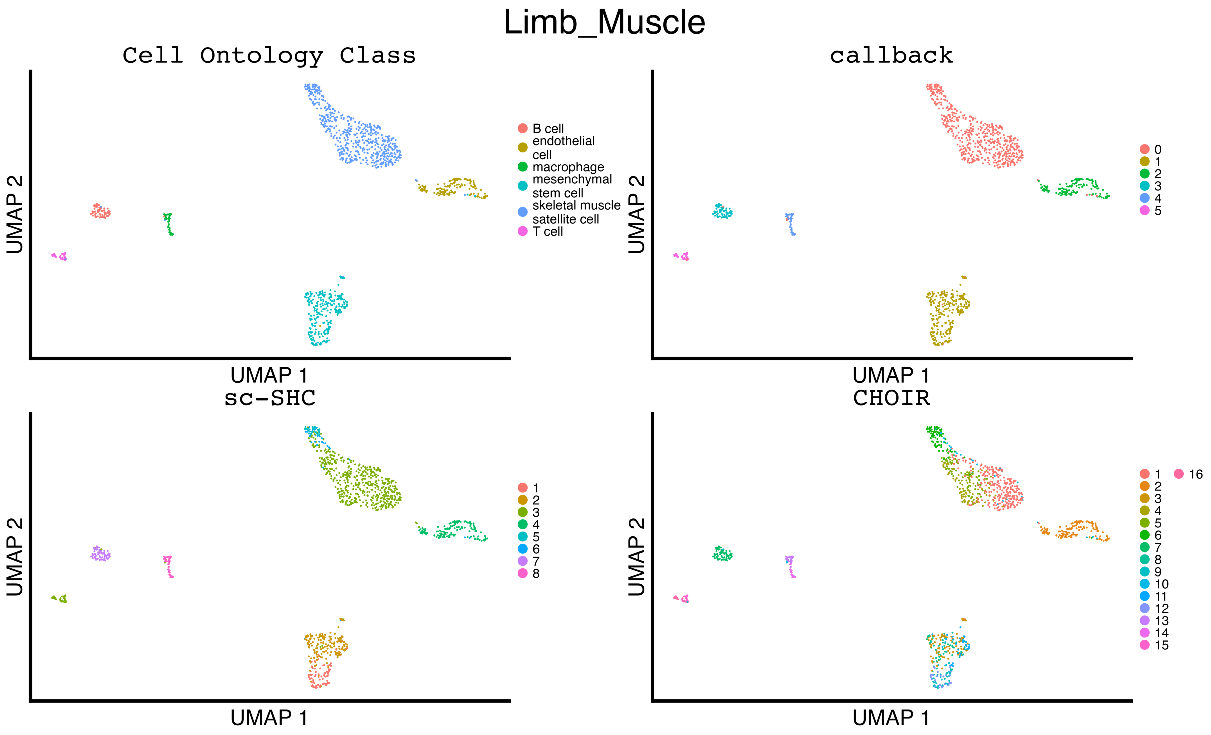


**Figure S15. Uniform manifold approximation and projection (UMAP) plots illustrating the manually curated cell ontology class labels compared to the inferred clustering results for `callback`, `sc-SHC`, and `CHOIR` when analyzing the limb muscle tissue from the Tabula Muris study.**

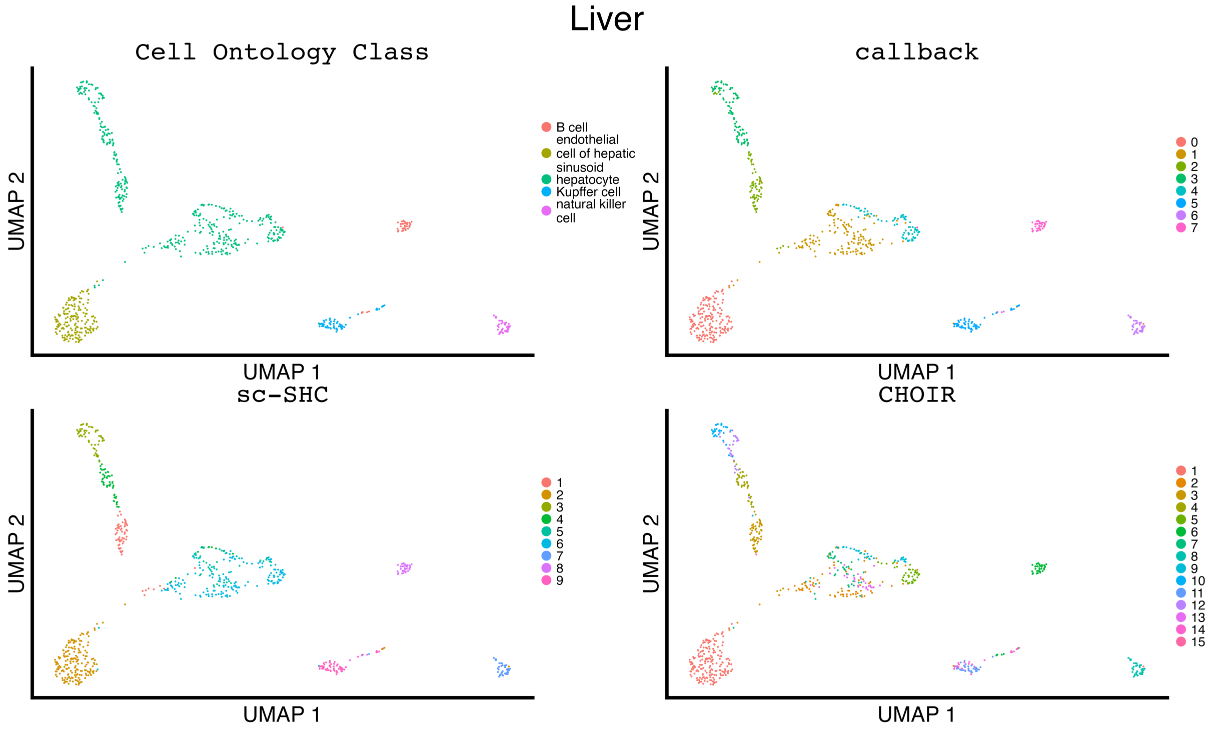


**Figure S16. Uniform manifold approximation and projection (UMAP) plots illustrating the manually curated cell ontology class labels compared to the inferred clustering results for `callback`, `sc-SHC`, and `CHOIR` when analyzing the liver tissue from the Tabula Muris study.**

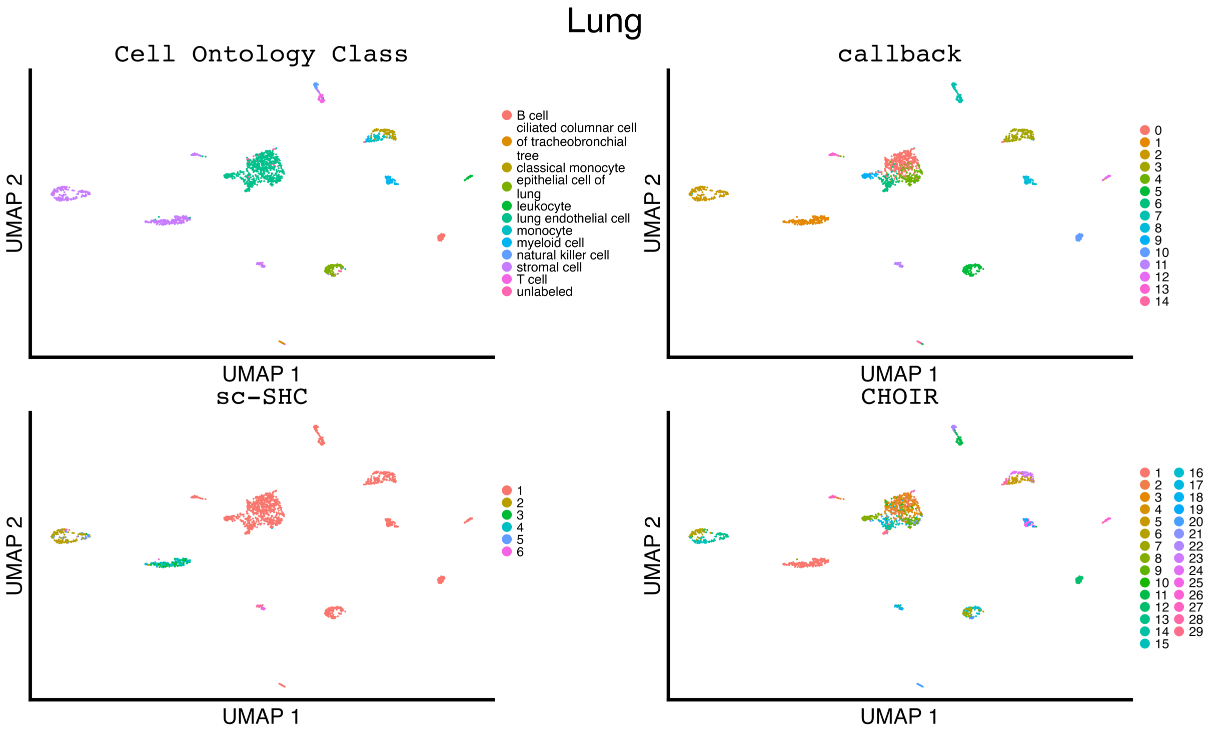


**Figure S17. Uniform manifold approximation and projection (UMAP) plots illustrating the manually curated cell ontology class labels compared to the inferred clustering results for `callback`, `sc-SHC`, and `CHOIR` when analyzing the lung tissue from the Tabula Muris study.**

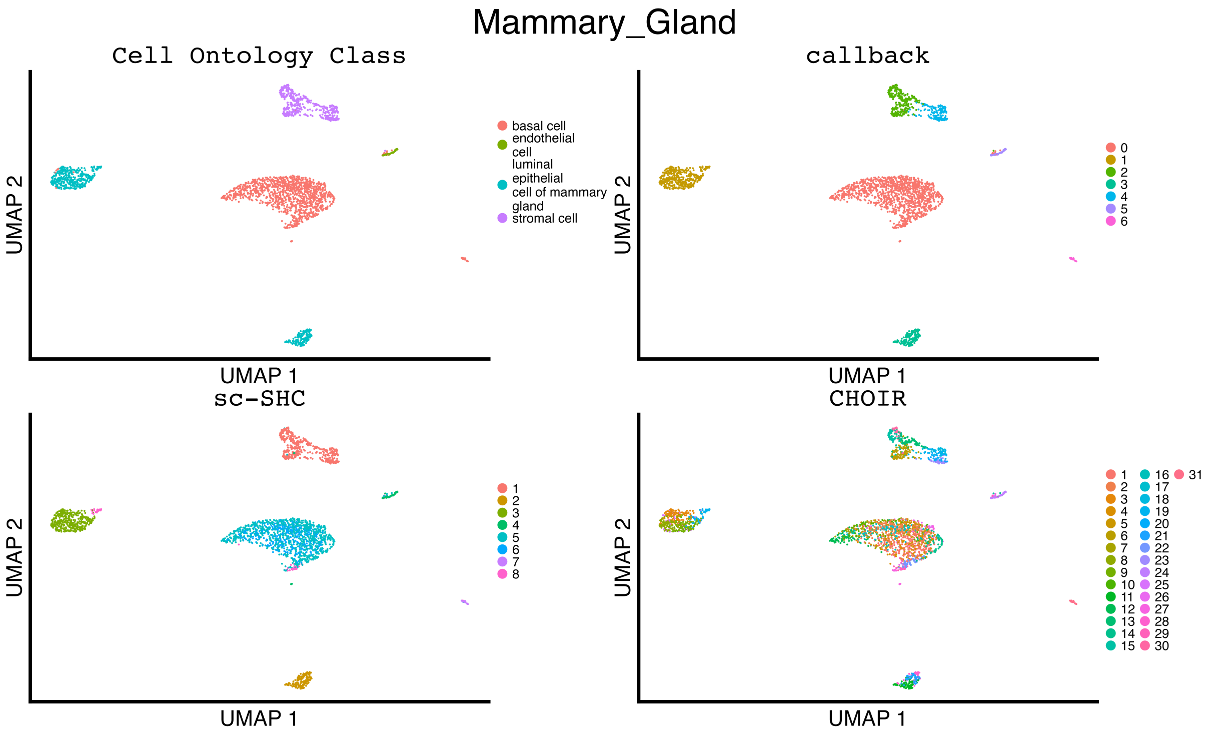


**Figure S18. Uniform manifold approximation and projection (UMAP) plots illustrating the manually curated cell ontology class labels compared to the inferred clustering results for `callback`, `sc-SHC`, and `CHOIR` when analyzing the mammary gland tissue from the Tabula Muris study.**

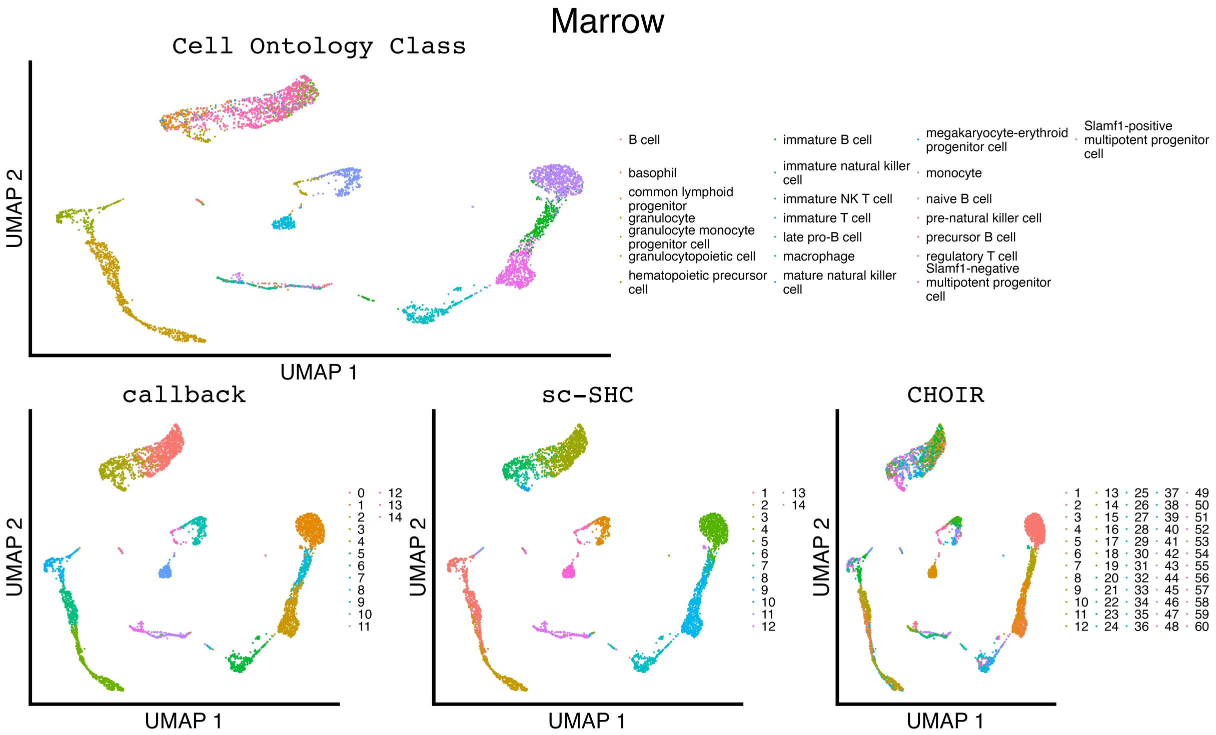


**Figure S19. Uniform manifold approximation and projection (UMAP) plots illustrating the manually curated cell ontology class labels compared to the inferred clustering results for `callback`, `sc-SHC`, and `CHOIR` when analyzing the bone marrow tissue from the Tabula Muris study.**

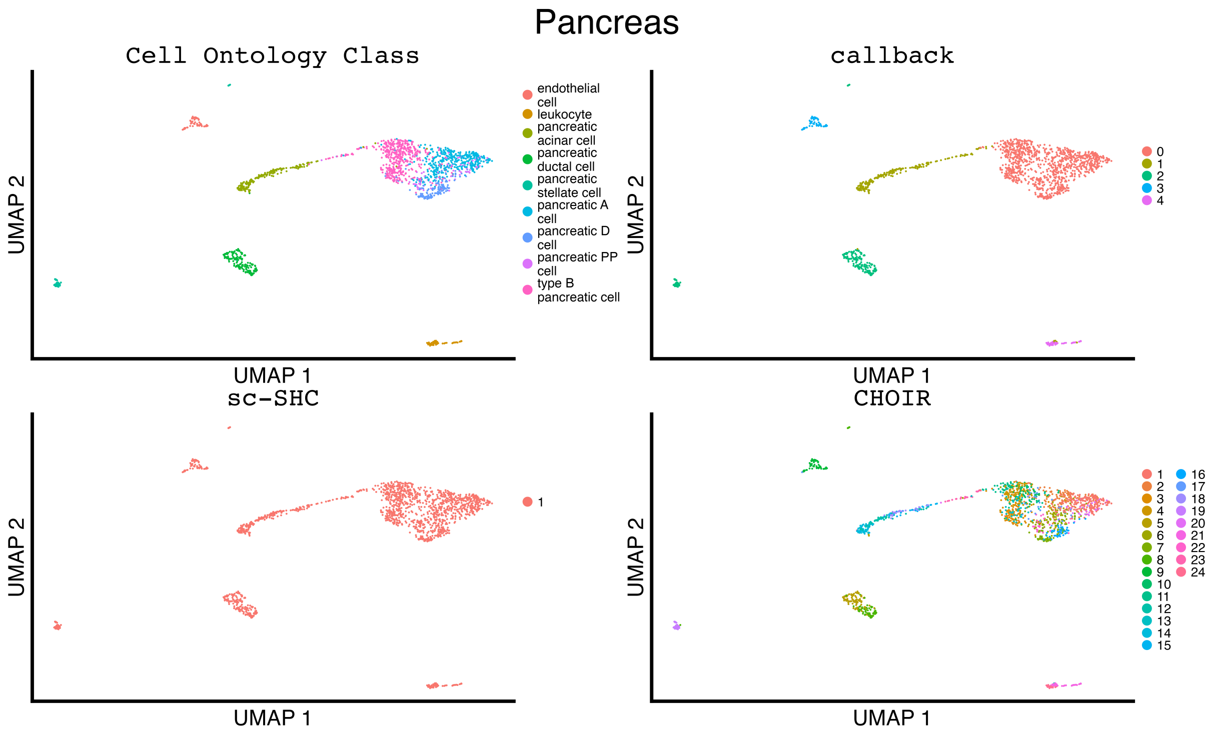


**Figure S20. Uniform manifold approximation and projection (UMAP) plots illustrating the manually curated cell ontology class labels compared to the inferred clustering results for `callback`, `sc-SHC`, and `CHOIR` when analyzing the pancreas tissue from the Tabula Muris study.**

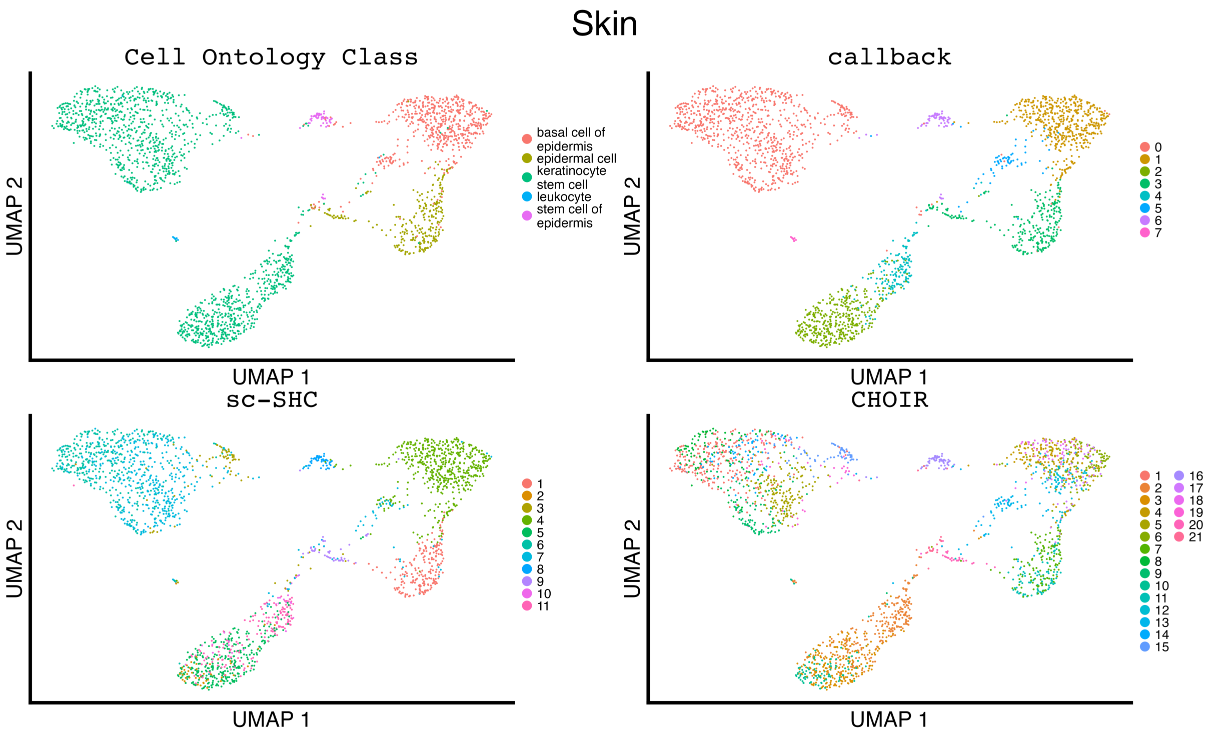


**Figure S21. Uniform manifold approximation and projection (UMAP) plots illustrating the manually curated cell ontology class labels compared to the inferred clustering results for** `callback`, `sc-SHC`, **and** `CHOIR` **when analyzing the skin tissue from the Tabula Muris study.**

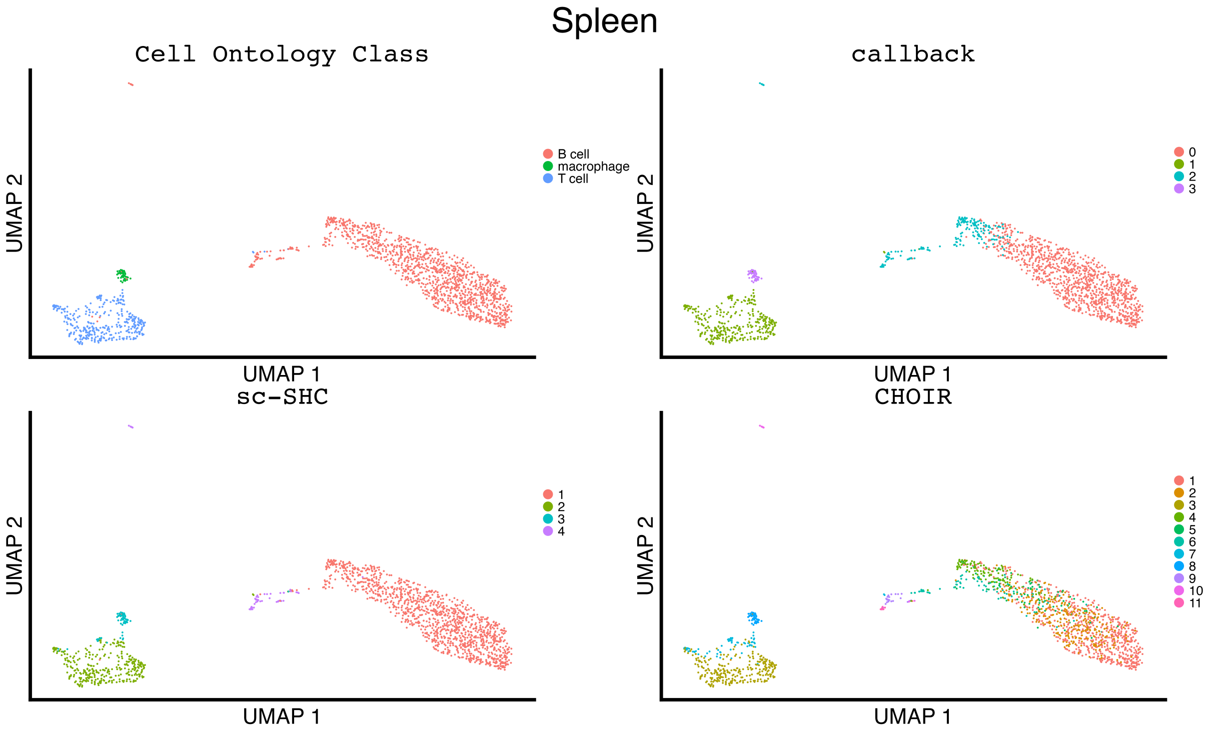


**Figure S22. Uniform manifold approximation and projection (UMAP) plots illustrating the manually curated cell ontology class labels compared to the inferred clustering results for `callback`, `sc-SHC`, and `CHOIR` when analyzing the spleen tissue from the Tabula Muris study.**

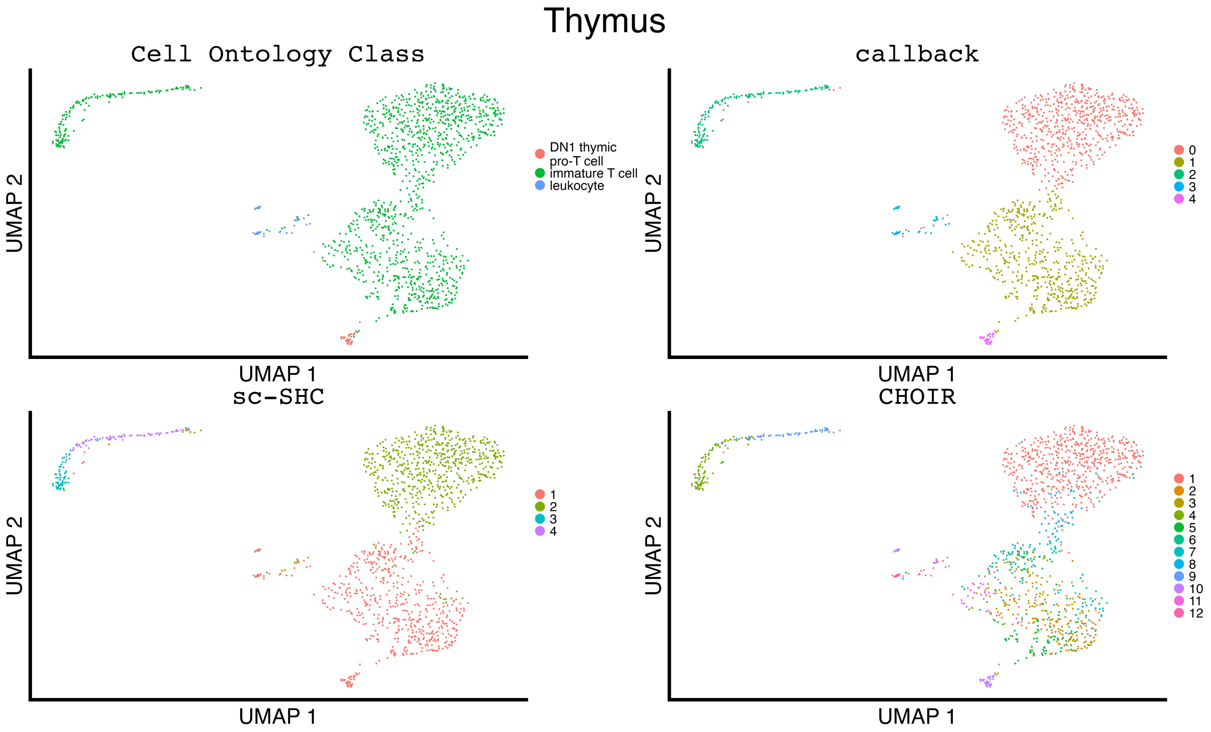


**Figure S23. Uniform manifold approximation and projection (UMAP) plots illustrating the manually curated cell ontology class labels compared to the inferred clustering results for** `callback`**,** `sc-SHC`**, and** `CHOIR` **when analyzing the thymus tissue from the Tabula Muris study.**

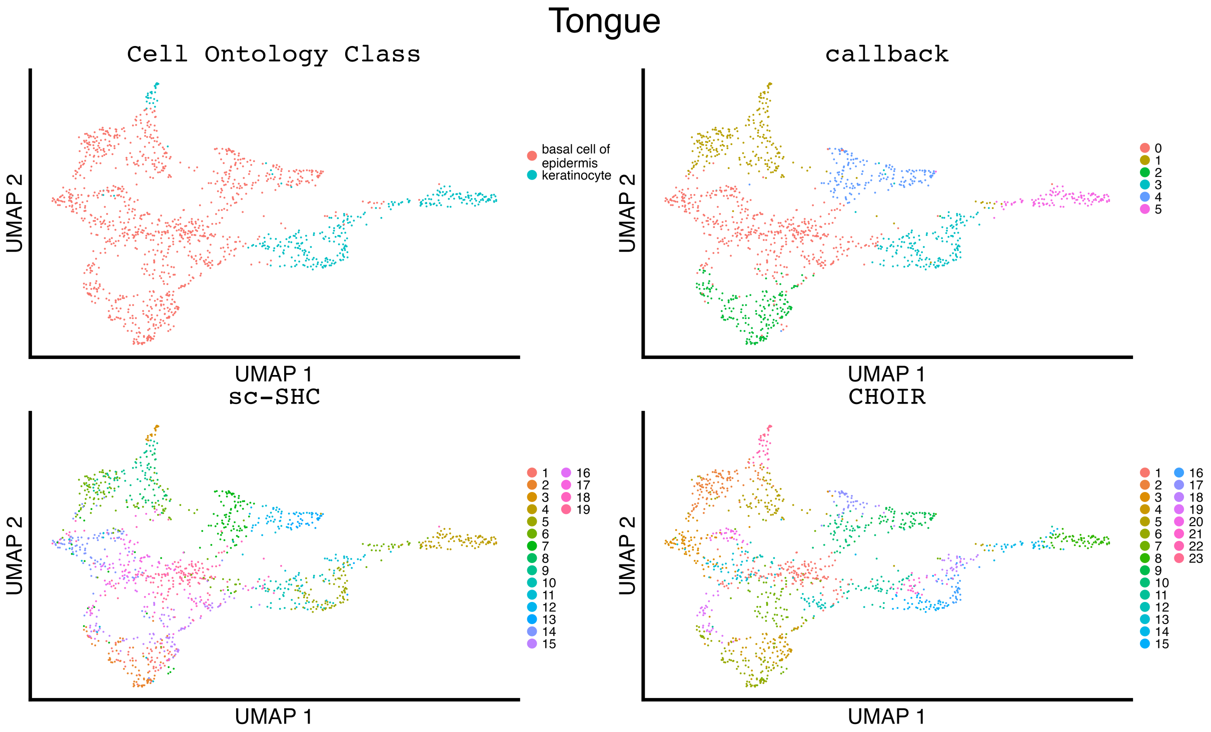


**Figure S24. Uniform manifold approximation and projection (UMAP) plots illustrating the manually curated cell ontology class labels compared to the inferred clustering results for `callback`, `sc-SHC`, and `CHOIR` when analyzing the tongue tissue from the Tabula Muris study.**

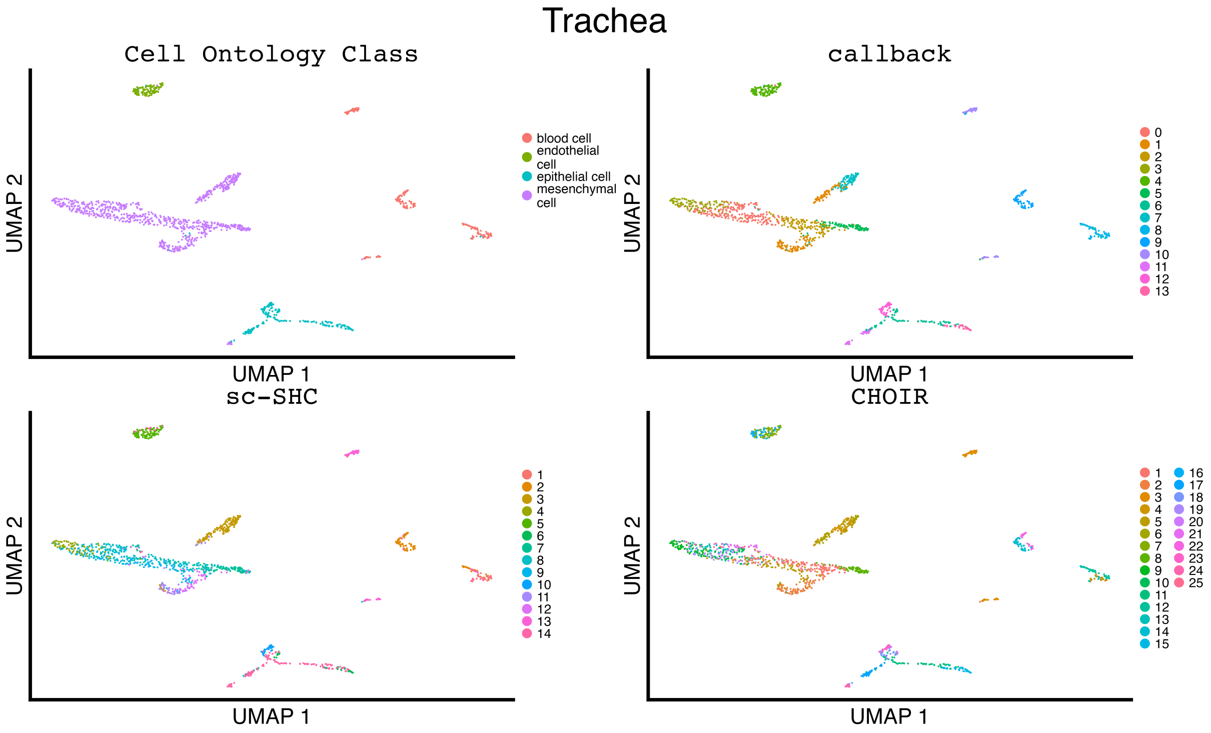


**Figure S25. Uniform manifold approximation and projection (UMAP) plots illustrating the manually curated cell ontology class labels compared to the inferred clustering results for `callback`, `sc-SHC`, and `CHOIR` when analyzing the trachea tissue from the Tabula Muris study.**

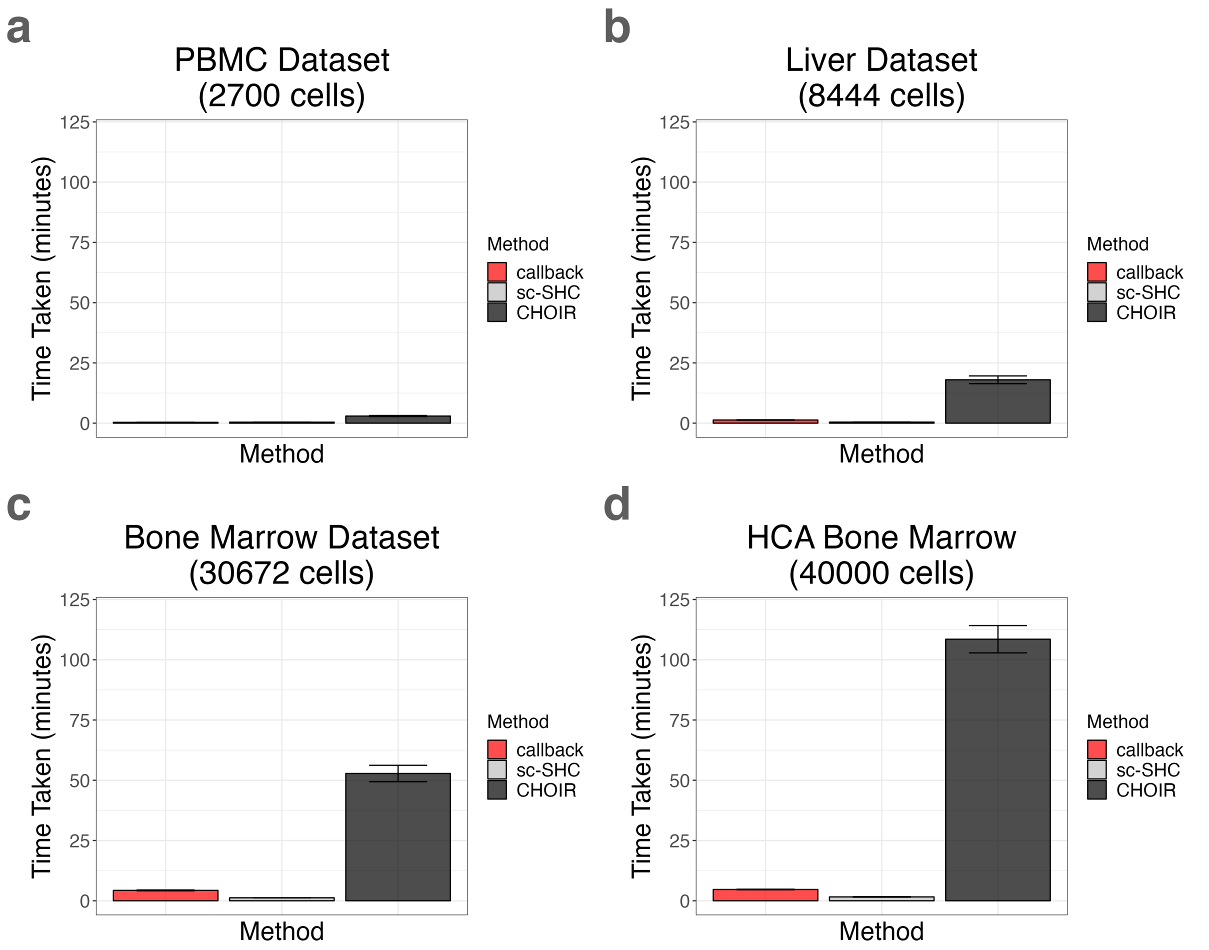


**Figure S26. Average runtime comparison of `callback`, `sc-SHC`, and `CHOIR` in four additional single-cell studies of varying sizes.** Each method was run on a machine with 16 cores. The datasets analyzed include **(a)** the PBMC 3K ($N$ = 2,700 cells), **(b)** the human liver data from MacParland et al. [6] ($N$ = 8,444 cells), and **(c, d)** the `SeuratData` bone marrow datasets ($N$ = 30,672 and $N$ = 40,000 cells, respectively). We run each method on each dataset 5 times; depicted in each bar plot is the mean $\pm$ the standard deviation across all runs.

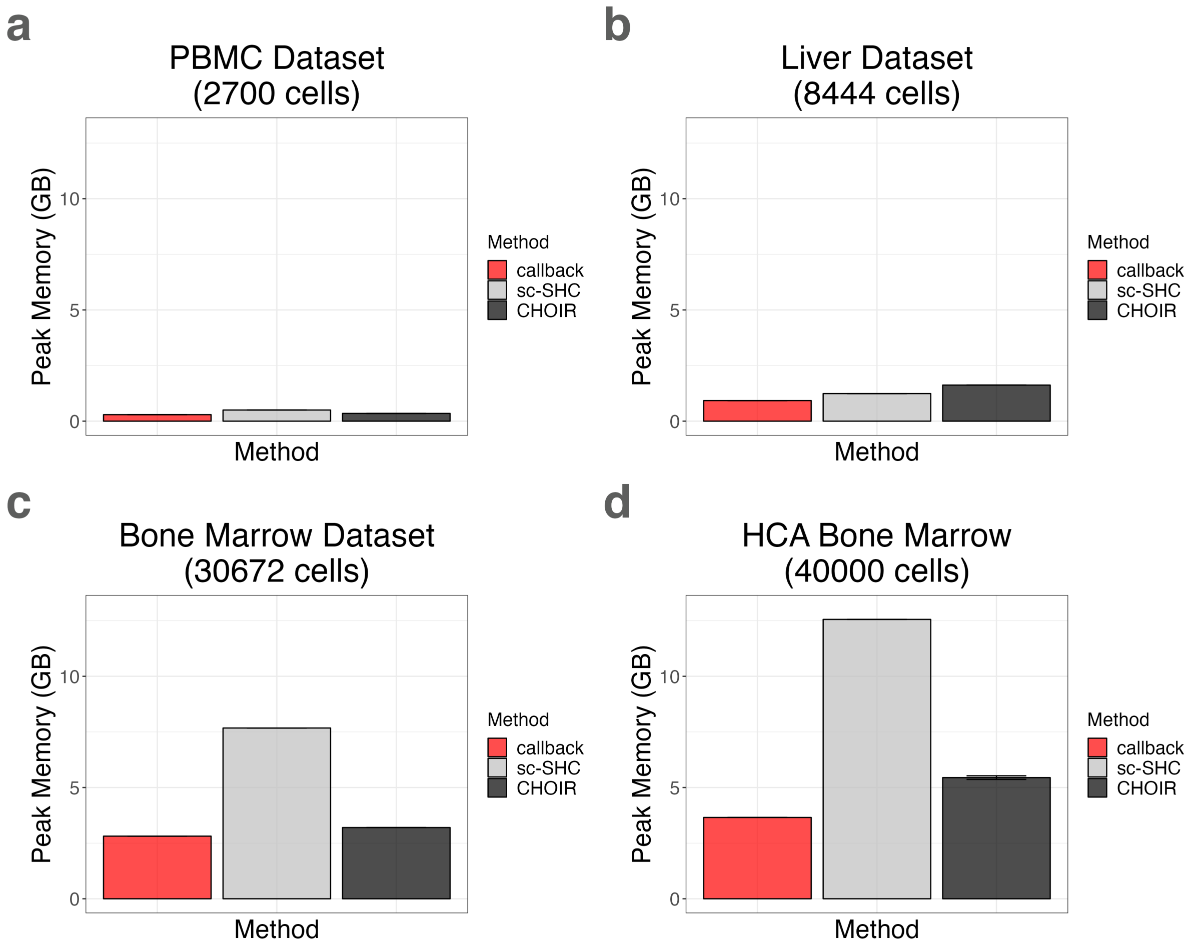


**Figure S27. Comparison of peak memory usage (in gigabytes; GB) for `callback`, `sc-SHC`, and `CHOIR` on four additional single-cell studies of varying sizes.** Each method was run on a machine with 16 cores. The datasets analyzed include **(a)** the PBMC 3K ($N$ = 2,700 cells), **(b)** the human liver data from MacParland et al. [6] ($N$ = 8,444 cells), and **(c, d)** the `SeuratData` bone marrow datasets ($N$ = 30,672 and $N$ = 40,000 cells, respectively). We run each method on each dataset 5 times; depicted in each bar plot is the mean ± the standard deviation across all runs.

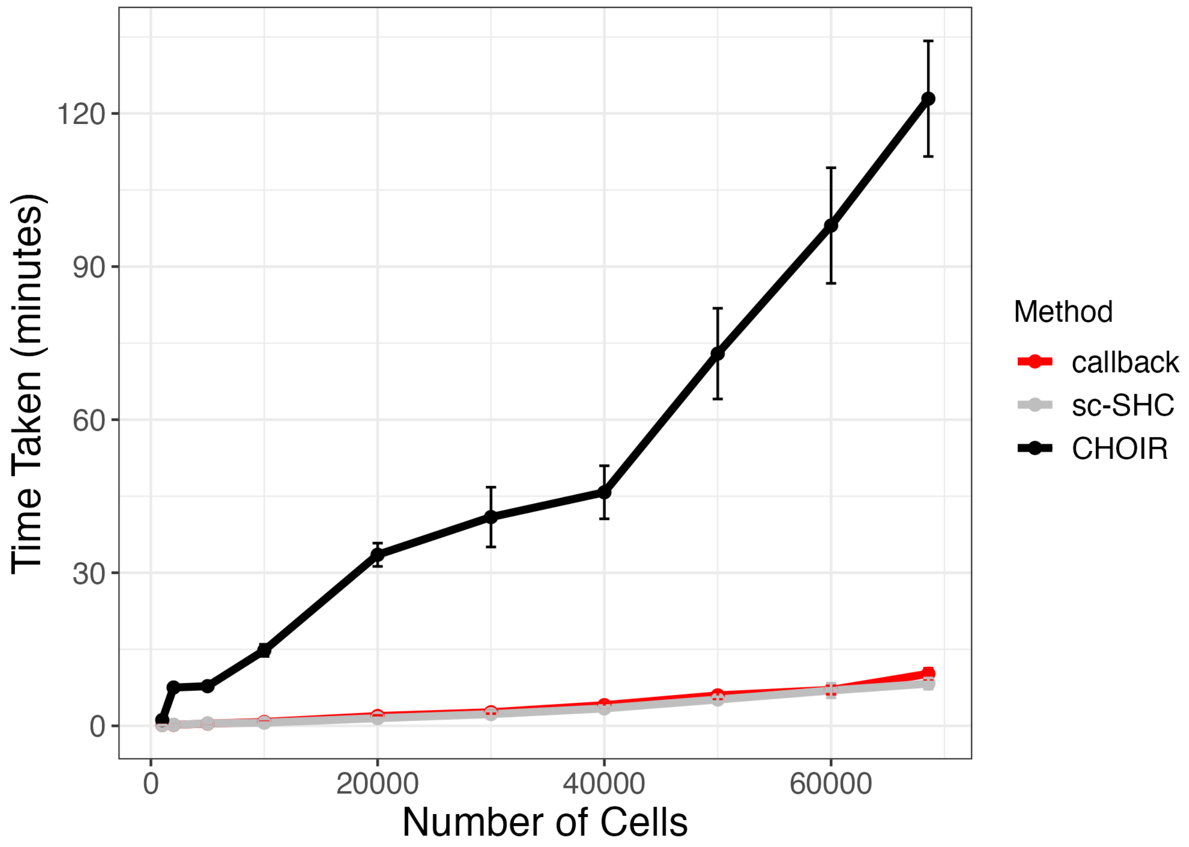


**Figure S28. Average runtime comparison of `callback`, `sc-SHC`, and `CHOIR` as a function of the number of cells in a study.** Here, we take subsets of the 68,579 total peripheral blood mononuclear cells (PBMCs) provided by Zheng et al. [7] which included smaller datasets of size 1K, 2K, 5K, 10K, 20K, 30K, 40K, 50K, and 60K cells. Each method was run on a machine with 16 cores. We run each method on each dataset 5 times; depicted are the mean ± the standard deviation across all runs.

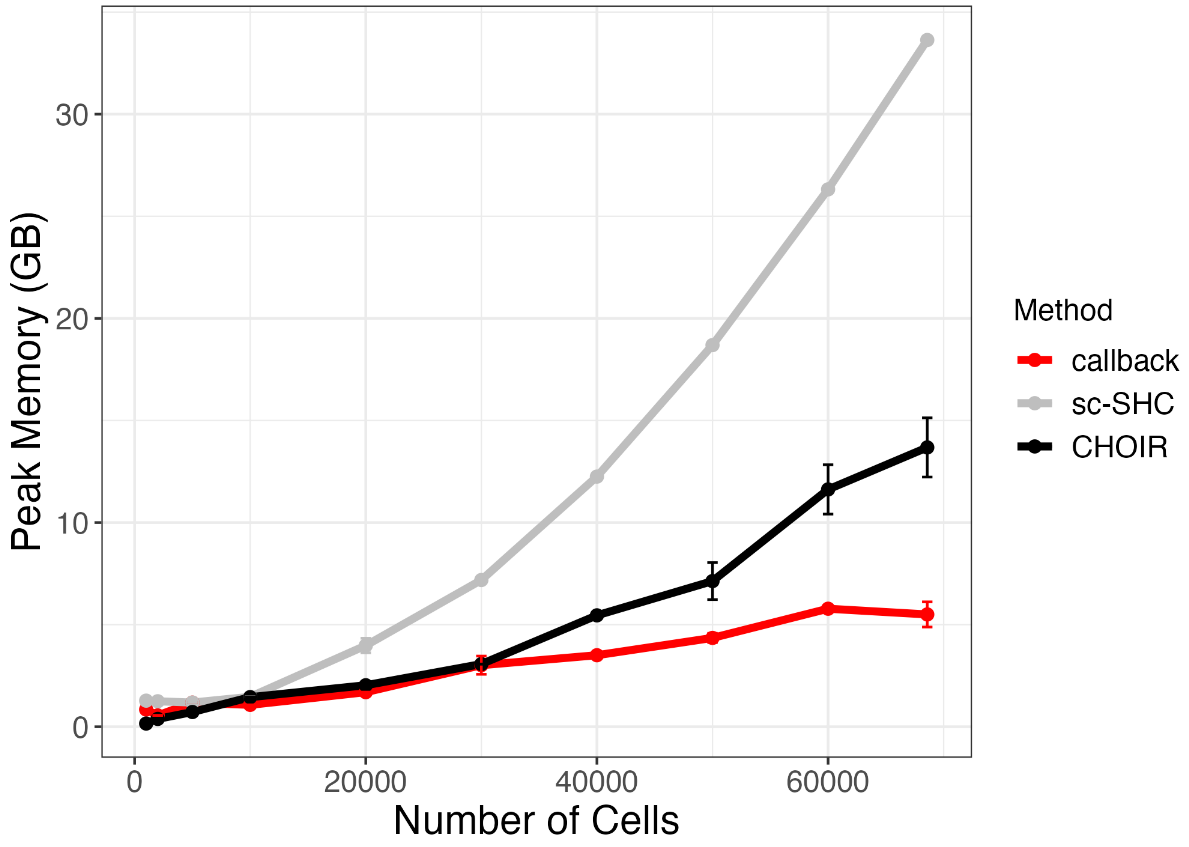


**Figure S29. Comparison of peak memory usage (in gigabytes; GB) for `callback`, `sc-SHC`, and `CHOIR` as a function of the number of cells in a study.** Here, we take subsets of the 68,579 total peripheral blood mononuclear cells (PBMCs) provided by Zheng et al. [7] which included smaller datasets of size 1K, 2K, 5K, 10K, 20K, 30K, 40K, 50K, and 60K cells. Each method was run on a machine with 16 cores. We run each method on each dataset 5 times; depicted are the mean ± the standard deviation across all runs.
